# The more, the merrier: multivariate phase synchronization methods excel pairwise ones in estimating functional brain connectivity from reconstructed neural sources

**DOI:** 10.1101/2023.01.19.524740

**Authors:** Ricardo Bruña, Pablo Cuesta, Federico Ramírez-Toraño, Isabel Suárez-Méndez, Ernesto Pereda

## Abstract

The estimation of functional connectivity (FC) from electro-(EEG) or magnetoencephalographic (MEG) recordings suffers from low spatial resolution, being one of the reasons for the reduced number of sensors compared to the number of reconstructed sources of activity. This problem can be avoided by estimating FC between larger regions instead of individual sources. However, combining all the sources in each area to produce a single time series per region is far from trivial.

We have used simultaneous EEG/MEG data from 11 participants and compared the FC estimates from both techniques by using different multivariate approaches. Since the underlying generators are identical for EEG and MEG, the more similar the FC estimation from both techniques is, the more accurate it is likely to be. The results show that using either the average or the root-mean-square of the bivariate source-to-source FC estimates consistently outperforms the use of a representative time series from each area.

We concluded that the reconstructed activity in each brain region is too complex to be reduced to a single representative time series and that full multivariate approaches must be used to describe between-area FC from electrophysiological recordings accurately. Moreover, the high correlation between the FC values estimated from EEG and MEG suggests that the results found in the high-sensitivity, low-noise MEG can be transferable to the more affordable EEG, at least when high-quality source reconstruction is used.

## 1. Introduction

For a long time, the scientific consensus was that the differentiated, highly specialized structures that comprise the brain worked separately to generate the brain function. However, the connectome (Sporns, Tononi, and Kötter, 2005) pictures the brain as a complex network of interconnected and highly specialized nodes (Friston, 1994).

Many brain diseases have been identified as disconnection syndromes in which not the brain areas themselves, but their communication is impaired (Geschwind, 1965; Mesulam, 2015). Consequently, functional neuroimaging has migrated from the study of isolated areas (Kwong *et al*., 1992) to brain connectivity (Biswal *et al*., 1995), often estimated by employing tools borrowed from the study of synchronization in complex systems. Synchronization can be operationally defined as a statistical relationship between some signals (e.g., their amplitudes or phases). Only when such dependence is higher than what would be expected by chance can we infer that their generating systems/processes can be connected. Thus, functional brain connectivity can only be studied as brain synchronization or, more precisely, synchronization in brain time-series activity.

Phase synchronization (PS) is a type of synchronization where the instantaneous phase of oscillatory processes is synchronized instead of its direct amplitude. PS was introduced by Rosenblum and colleagues in 1996 (Rosenblum, Pikovsky and Kurths, 1996) and was quickly accepted as a synchronization model for the brain (Lachaux *et al*., 1999; Mormann *et al*., 2000). One crucial point in favor of PS as a brain synchronization model is that the phase lock can be established with minimal energy. In contrast, an amplitude lock requires a more significant amount of that energy and reduces the degrees of freedom of the systems involved.

When studying functional synchronization in the brain and parting from the idea that all the brain activity is meaningful to our experiment, we need to consider all possible interactions between all potential activity sources. Neurophysiological techniques such as electroencephalography (EEG) or magnetoencephalography (MEG) record the signals extracranially and, therefore, the activity sources must be reconstructed by using an inverse method. The number of those reconstructed sources ranges between the thousands and the tens of thousands, depending on the employed inverse method.

For MEG and EEG, signals are acquired simultaneously from hundreds of sensors/electrodes. As a result, close brain sources are highly correlated, and the information is redundant. To solve this problem, neighboring sources are grouped into areas, a process named parcellation, which, at the same time, reduces the amount of information in the problem. These areas are usually defined using anatomical atlases (Tzourio-Mazoyer *et al*., 2002; Desikan *et al*., 2006). After this grouping, whole-brain synchronization can be measured as synchronization between pairs of areas instead of pairs of sources. The use of these atlases carries an extra advantage: the number of regions in the atlas is the same for all participants, and, thereby, it makes the comparison of results easier.

When using a region-based approach to evaluate brain synchronization, the algorithms must be applied to pairs of multivariate sets of signals (i.e., the sources in each region). Since synchronization algorithms are typically developed to calculate pairwise synchronization, their original formulation cannot be directly used. One particular solution can be reducing the dimensionality of the data by combining the signals from all the sources in each brain area to generate a single time series representative of the area, which comprises as much information of the original data as possible (Korhonen, Palva and Palva, 2014). Popular approaches include, but are not limited to: the use of a single source, placed on the centroid of the brain area (Luckhoo *et al*., 2012); the use of the source whose time series is more correlated to the rest of the sources in the area (Garcés, Martín-Buro, and Maestú, 2016; López-Sanz *et al*., 2017); the use of the component, as calculated with PCA, that explains the largest amount of variance of the whole area (Friston *et al*., 2006; King *et al*., 2015; Schwabedal and Kantz, 2016); or the average of the time series in the area (Craddock *et al*., 2012; James, Hazaroglu and Bush, 2016).

An allegedly more principled approach would imply the use of synchronization metrics expanded to the case of multivariate data, i.e., to tackle the particular instance of estimating synchronization between two sets of *N* and *M* time series (*N, M* >1; being *N* not necessarily equal to *M*). For example, the famous correlation coefficient can be expanded through canonical correlation (Hotelling, 1936). A similar approach can be used to extend the corrected imaginary part of coherence (Pascual-Marqui, 2007b) to a multivariate scenario (Ewald *et al*., 2012).

A third approach, also a multivariate one, makes use of the algebraic properties of the *NxM*, pairwise synchronization matrix. This method has already been used, for example, to extend the Phase Slope Index (Nolte *et al*., 2008) into its multivariate counterpart (Basti *et al*., 2018).

In this study, we have evaluated different methods that extend inter-area PS estimation to a multivariate scenario. Note that we cannot directly determine each technique’s validity by comparing its estimation to a gold standard because the actual brain FC pattern is unknown in an experimental situation. Instead, we assess its quality through an indirect process, by considering that EEG and MEG are two different manifestations of the same electrophysiological phenomenon and describe the same underlying FC. Thus, a reasonable estimation of this otherwise unknown FC should render similar results for both techniques. Consequently, by starting with a set of simultaneous EEG and MEG recordings and comparing their inter-area PS estimated using these different methods, we will regard the one producing the largest resemblance between the EEG- and the MEG-derived FC estimates.

We hypothesize that the richness of the information included in the model will be paramount, and using the information contained in all the sources in each area will produce better results than using any representative time series. This hypothesis is based on the idea that the anatomical limits will not necessarily match the functional limits, and each parcellation area will likely contain several units of information. Using one representative time series per area will inevitably entail losing information, selecting only one of the information units, and disregarding the rest. In our particular case, the representative time series approaches will likely select different features for EEG and MEG, thereby giving different results. Instead, using all the information in each area will ensure that the amount of information coming from both techniques is roughly the same. For the same reason, these results are also expected to be closer to the ground truth.

## 2. Materials and methods

### 2.1. Participants

Eleven participants (age 77.9 ± 6.4, mean ± standard deviation, eight women) were enrolled in this project. All participants signed informed consent before enrollment. This study was approved by the “Hospital Universitario San Carlos” ethics committee, and the procedures were performed following approved guidelines and regulations. The data and code used in this work are available under the terms listed in section 6 Data and code availability.

We acquired simultaneous EEG and MEG data using an Elekta Vectorview system (Elekta, AB, Stockholm, Sweden) placed inside a magnetically shielded room (VacuumSchmelze GmbH, Hanau, Germany) at the Center for Biomedical Technology in Madrid (Spain). MEG data were acquired using the system’s 306 sensors (102 magnetometers, 204 planar gradiometers). EEG data were obtained using 32 electrodes EasyCap (Brain Products GmbH, München, Germany) referenced to both earlobes’ average. Besides, we used two pairs of bipolar electrodes to capture the electrooculographic and electrocardiographic artifacts. The montage was completed with four head position indication (HPI) coils, two on the forehead and two on the mastoids. The anatomical landmarks at the preauricular points and the nasion, the HPI coils position, and the electrode positions were acquired using a 3D FASTRAK digitizer (Polhemus, Colchester, Vermont). Also, we recorded over 200 head shape points to aid in the realignment of the head model.

### 2.2. Signal acquisition and pre-processing

We acquired two sessions of 5 minutes of resting-state data with eyes closed for each participant. The acquisition set up was the same for MEG and EEG, using an anti-alias band-pass filter between 0.1 and 330 Hz and a sampling rate of 1000 Hz. MEG data were pre-processed using the temporal extension of the signal space separation (tSSS) method (Taulu and Simola, 2006), using 10 seconds as window length and 0.9 as correlation limit. Continuous head position was active during the recording and used to off-line correct head movements of the participants.

Data underwent automatic artifact selection using FieldTrip (Oostenveld *et al*., 2011), and a MEG expert confirmed the findings. After the artifact’s removal, we applied second-order blind identification (SOBI) (Belouchrani *et al*., 1997) separately for EEG and MEG to remove cardiographic, oculographic, and noise-related components. Only the magnetometers’ data were used for MEG since, after tSSS, the sensor-space data is highly redundant (Garcés *et al*., 2017). The resulting time series were segmented in epochs of 4 seconds of artifact-free data. Three were discarded from the original 22 sessions due to high noise, resulting in 19 sessions with 63.5 ± 9.5 epochs each (minimum of 42). Data are described in Table 1.

**Table 1.**
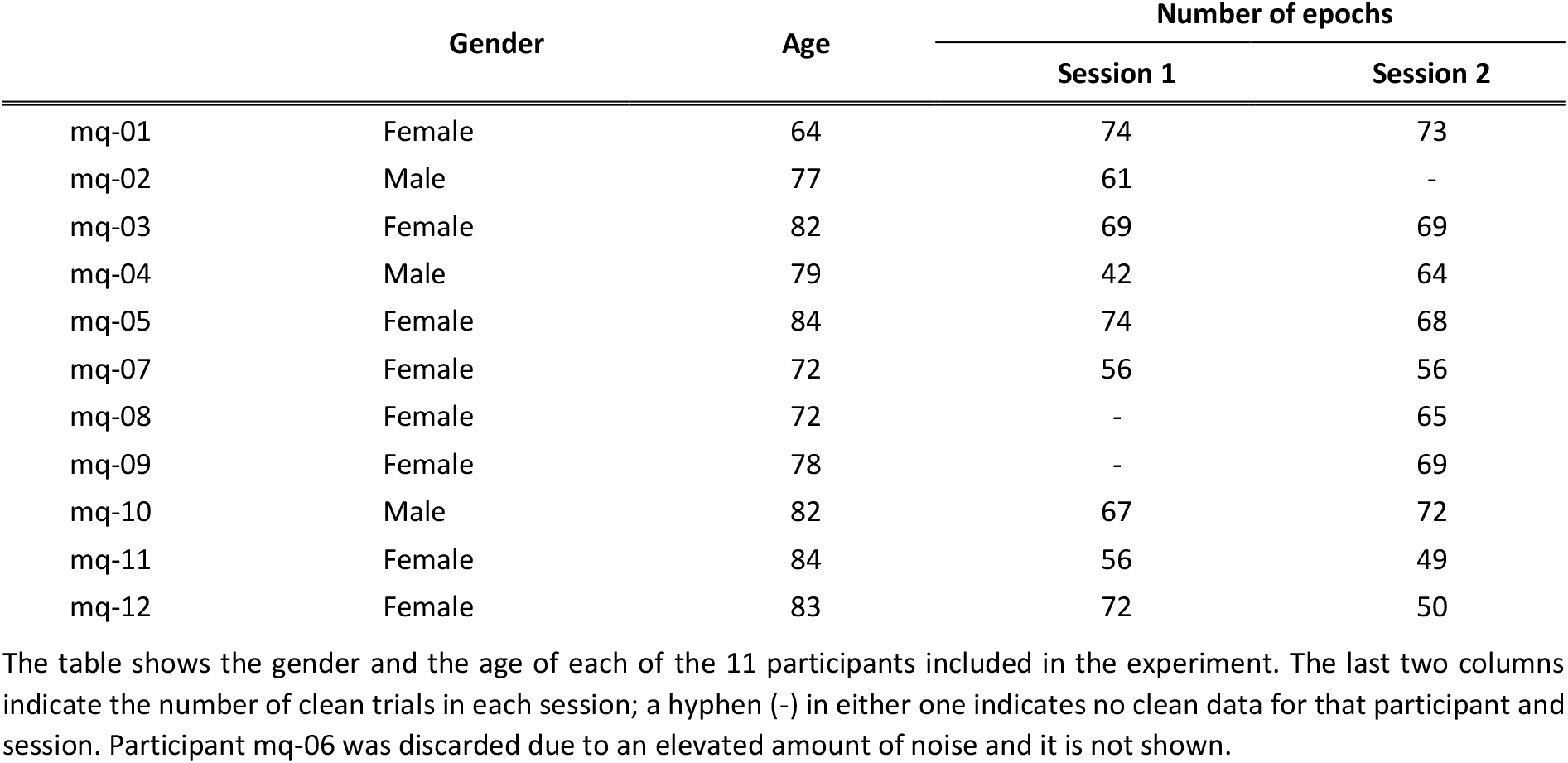
Sociodemographic data of the participants.

|  | Gender | Age | Number of epochs |  |
| --- | --- | --- | --- | --- |
|  |  |  | Session 1 | Session 2 |
| mq-01 | Female | 64 | 74 | 73 |
| mq-02 | Male | 77 | 61 | - |
| mq-03 | Female | 82 | 69 | 69 |
| mq-04 | Male | 79 | 42 | 64 |
| mq-05 | Female | 84 | 74 | 68 |
| mq-07 | Female | 72 | 56 | 56 |
| mq-08 | Female | 72 | - | 65 |
| mq-09 | Female | 78 | - | 69 |
| mq-10 | Male | 82 | 67 | 72 |
| mq-11 | Female | 84 | 56 | 49 |
| mq-12 | Female | 83 | 72 | 50 |

The table shows the gender and the age of each of the 11 participants included in the experiment. The last two columns indicate the number of clean trials in each session; a hyphen (-) in either one indicates no clean data for that participant and session. Participant mq-06 was discarded due to an elevated amount of noise and it is not shown.

### 2.3. Source reconstruction

All the participants had anatomical images (T1 MRI or CT), which we segmented using the *New segmentation* algorithm in SPM12 (Ashburner and Friston, 2005) on the brain (understood as the union of gray matter, white matter, and cerebrospinal fluid), bone, and soft tissue. We used this segmentation to create a set of surface interfaces with iso2mesh (Fang and Boas, 2009), and then a realistic boundary element method (BEM) head model using OpenMEEG (Gramfort *et al*., 2010). As a source model, we used a homogeneous three-dimensional grid with a spacing of 10 mm, defined in MNI space. Each source position was labeled using the Automated Anatomical Labeling (AAL) atlas (Tzourio-Mazoyer *et al*., 2002). Only those sources labeled as part of one of the 80 cortical areas of the atlas were used. Finally, this grid was linearly transformed to subject space using the individual anatomical image, resulting in a subject-specific source model. The subject-specific head and source models were realigned to MEG space using the anatomical landmarks and the digitized head shape as a guide and then employed to create individual lead fields for EEG and MEG, which were used as forward models.

As an inverse solution, we used a linearly constrained minimum variance (LCMV) beamformer (van Veen *et al*., 1997), as it has been proven to behave appropriately for the estimation of resting-state functional connectivity (Hincapié *et al*., 2017). The spatial filter was built separately for EEG and MEG, using the subject-specific, trial-averaged covariance matrix, and a regularization factor of 10 % for the average sensor power. Following the literature’s usual approach, we calculated a scalar beamformer solution by projecting the primary orientation vector solution. This procedure results in a single time series per source position. To maximize the source-reconstructed data, we also generated a vector beamformer solution, namely a 3D beamformer, containing 99 % of the information and not only the primary orientation. We thoroughly detail the construction of this vector beamformer in the Supplementary materials. In the Figures, Tables, and text, the indices calculated using this approach are marked with a “3D” superscript.

### 2.4. Phase synchronization

We evaluate PS using the phase-locking value (PLV) (Mormann *et al*., 2000). PLV is a bivariate PS metric, ranging between zero and one, which quantifies the degree of phase-locking of two-time series. Mathematically, PLV is calculated as follows:

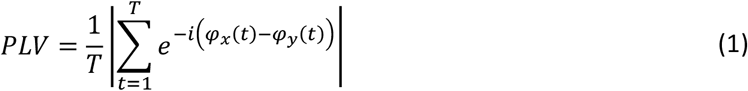

where *T* is the length of the signals in samples, and *φ*_*x*_ (respectively, *φ*_*y*_) is the instantaneous phase of signal *x* (respectively, signal *y*). For the value of PLV to make sense, *x* and *y* must be narrowband signals originated by an oscillatory process. In this work, we obtained the signal’s instantaneous phase through the Hilbert transform of the narrowband signal, using 2 seconds of real data as padding to avoid edge effects.

The selection of PLV as the metric of interest is based on its extensive use and the existence of an optimized definition (Bruña, Maestú, and Pereda, 2018), which allows for the calculation of whole-brain connectivity in a few minutes in an average computer. While popular, PLV has some well-known caveats. Firstly, it is susceptible to volume conduction effects, thereby unable to tell apart a source leakage’s effects from a genuine source interaction. Secondly, the metric definition, based only on the narrowband signal phase, is nonlinear, and this might introduce some instability issues, for example, when dealing with spatial rotations (Ewald *et al*., 2012).

To overcome the first limitation, we also studied a derived metric, the corrected imaginary PLV (ciPLV) (Bruña, Maestú, and Pereda, 2018), insensitive to instantaneous interactions, effectively removing the effect of source leakage in doing so. Its definition is similar to that of PLV:

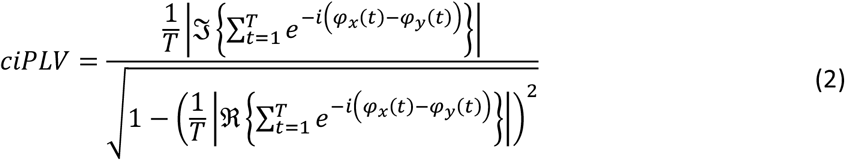

where ℛ{•} and 𝒥{•} stand for the real and imaginary parts, respectively.

To deal with the second limitation, we used a linear extension of PLV, the Hilbert coherence (HCoh) (Bruña, Maestú, and Pereda, 2018). This metric’s main advantage is that, since it is linear, it is transparent to linear source reconstruction algorithms (such as beamformer) and can be efficiently computed in sensor space and then transformed to source space (Gross *et al*., 2001). The definition of HCoh closely recalls that of coherence (Nunez *et al*., 1997):

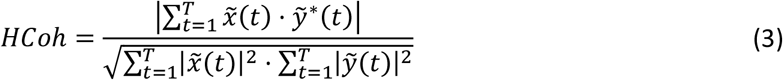

where 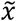 contains both the phase and the amplitude of *x*, and 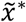 is its complex conjugate. 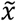 is obtained using Hilbert’s analytical signal, thus the name of the metric. Additionally, source leakage can be effectively removed from HCoh using a similar transformation to that shown in (2), obtaining the corrected imaginary Hilbert coherence (ciHCoh).

### 2.5. Multivariate approaches

In this study, we will evaluate the ability to extract genuine inter-area PS of several classical approaches in order to define one representative time series from each area: the source closest to the centroid of the brain area; the dominant component in the brain area, using principal component analysis (PCA); and the average of all the time series in the brain area. Besides, we will evaluate two purely multivariate approaches based on the synchronization matrix. The first one will be the average of all the bivariate PS values, i.e., the inter-area synchronization matrix average. The second one will be an extension of the cluster analysis for multivariate data to inter-area synchronization.

Some multivariate PS metrics have been described (Allefeld and Kurths, 2004; Al-Khassaweneh *et al*., 2016) to extract a global synchronization index. The idea is to apply cluster analysis to the pairwise PS matrix, identifying both the clusters themselves and each source’s participation coefficient in each cluster (Allefeld and Kurths, 2004). This idea can be achieved by utilizing eigen-decomposition of the pairwise PS matrix, where the eigenvectors indicate the participation coefficients of each source in each cluster, and the corresponding eigenvalues indicate the global variance explained by the cluster (Lohmann *et al*., 2010; Groth and Ghil, 2011). A global measure can be obtained from the root-mean-squared of the normalized eigenvalues:

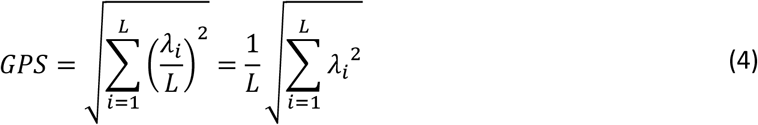

where *GPS* is a global PS index, and *λ*_*i*_ is the *i*-th eigenvalue of the *L* by *L* pairwise PS matrix. From this definition, GPS equates to the normalized Frobenius norm of the pairwise PS matrix.

Another approach uses hyperdimensional geometry to consider all the pairwise PS values in the matrix (Al-Khassaweneh *et al*., 2016). By defining the vector of phase differences in hyperspherical coordinates rather than polar ones, we can obtain a global synchronization index, named hyperspherical PS (HPS):

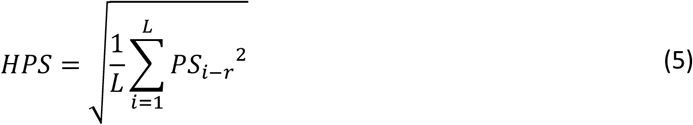

where *PS*_*i*−*r*_ is the bivariate PS calculated between time series *i* and a reference time series *r*. The selection of the reference oscillator plays an essential role in the stability of HPS. To avoid this issue, we can replace the original definition of HPS with the average HPS using different time series as oscillators:

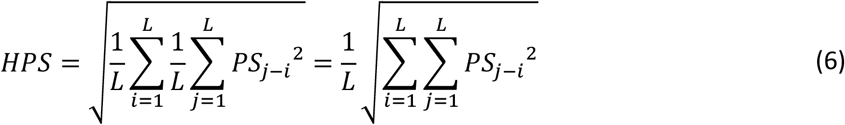

That is, our re-defined HPS is equal to the root-mean-square of the pairwise PS values. Once again, this definition equates to the normalized Frobenius norm of the pairwise PS matrix, and GPS and HPS are thus equivalent.

As the Frobenius norm can be calculated for rectangular nonsymmetrical matrices, it is possible to extend the GPS/HPS definition to get an index of inter-area PS:

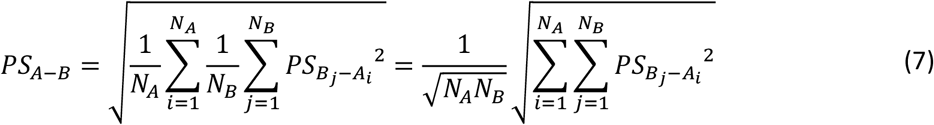

where *N*_*A*_ is the number of sources in area *A*, and *A*_*i*_ is the *i*-th source of that area. We termed this approach the root-mean-square (RMS) of the pairwise PS, and it is the last approach we will use to estimate inter-area PS.

It is noteworthy that, while we refer to these metrics as multivariate, an alternative, perhaps more accurate nomenclature would be mass-bivariate approaches. Nevertheless, similar metrics are termed multivariate in the literature (Allefeld and Kurths, 2004; Ewald *et al*., 2012; Al-Khassaweneh *et al*., 2016; Basti *et al*., 2018). Thus, we have decided to use this most straightforward, even if not entirely accurate, convention (see the Supplementary Materials for a thorough discussion on the topic).

### 2.6. Statistical analysis

Even when generated by the same underlying activity, EEG and MEG are inherently different. EEG signals are more affected by the skull’s low conductivity, which acts as a spatial, low-pass filter. On the other hand, MEG is insensitive to one source orientation (the radial orientation in spherical models), thereby missing some information about the sources’ activity. With this in mind, it would be unrealistic to expect identical results in the FC estimated from both techniques. We do expect, however, to find similar variations in the estimated FC values; that is, if one participant shows an increased (meaning, with respect to the mean of the distribution of all FC values) FC value in one connection between two brain areas as measured by EEG, we expect a similar behavior, as measured by MEG, between those same areas. We quantify the similitude of the FC values derived from EEG and MEG using a simple connection-wise correlation. For each link connecting two areas *i* and *j*, we calculate the Pearson correlation coefficient, for the 19 measurements, between the value of the link as estimated from EEG data, and the value estimated from MEG data. As we expect a positive relation between EEG and MEG, the correlation was evaluated one-tailed, considering only the positive values.

Since the number of comparisons is large (3160 inter-area FC plus 80 intra-area values), we expect many statistically significant correlations to be found just by chance. To handle this “multiple comparisons” problem, we have used two distinct approaches. First, we performed an ad-hoc correction by looking at the ratio of significant correlation values, in addition to the values themselves. If the FC estimated from EEG and MEG were completely independent, we would expect, by chance, a 5 % of significant correlations (or 2.5 % of significant positive correlations), with an alpha level of 0.05, in the complete pool of data. As we expect a relation between EEG and MEG (both are estimators of the same ground truth), the percentage of significant correlations in our data should be much higher. In fact, the closer the estimation of FC from EEG and MEG is to the ground truth, the larger this percentage should be. Secondly, we applied the idea of a false discovery rate (FDR) (Benjamini and Hochberg, 1995). This framework estimates the number of significant results we would expect in a completely random sample, and dynamically adjusts the significance threshold to remove them. Consequently, for a random set of data, the number of significant results, after correction, should be zero. Thus, we will evaluate the different approaches using the rate of significant (positive) correlations between EEG and MEG.

## 3. Results

An estimator of PS can be deemed as a valid FC metric when a meaningful phase can be derived from the signals so that they can, indeed, synchronize. In this work, we used Hilbert transform to estimate the instantaneous phase of our signals, and for this phase to have a physical meaning, we calculated it over narrowband versions of the signals. Namely, we calculated the PS band-wise, using the classical definition of the electrophysiological bands: theta (from 4 to 8 Hz), alpha (from 8 to 12 Hz), low beta (from 12 to 20 Hz), high beta (from 20 to 30 Hz), and gamma (from 30 to 45 Hz). The results for all the studied bands are shown in the supplementary materials. In this section, we only show the results for bands theta, alpha, and low beta. The rationale behind this comes from the very definition of PS: PS requires a meaningful phase, and while we would expect oscillators in theta and alpha, it is not clear if there exists an oscillator in low beta, so we evaluated both cases.

Figures 1 to 4 (along with Supplementary Figures 3 to 7) show the results using violin plots. We think it may be helpful to provide here some hints on how to interpret these results. These types of plots are a useful tool to represent probability distributions. Similar to box plots, violin plots allow for comparing different distributions but add extra information on their shape. In this case, we show distributions of correlation coefficients, ranging from -1 to 1, and depict the similitude between PS calculated from EEG and MEG across the 19 sessions considered. Therefore, each plot is composed of the 3160 links connecting each possible pair of areas. Besides the shape of the probability distribution, the violins represent the median (marked as a strong point representing the 50^th^ percentile) and the interquartile range (with the 25^th^ and 75^th^ percentile marked as light points connected with the median by a line). Finally, a dashed line in each figure marks the correlation level as significant (only positive correlation with an alpha level of 0.05) for a distribution of 19 pairs of elements. In a random distribution, this dashed line should define approximately 5 % of the distribution. Deviations from this number can be interpreted as deviations from a random distribution, and therefore as results not explainable solely by chance.

**Figure 1.**
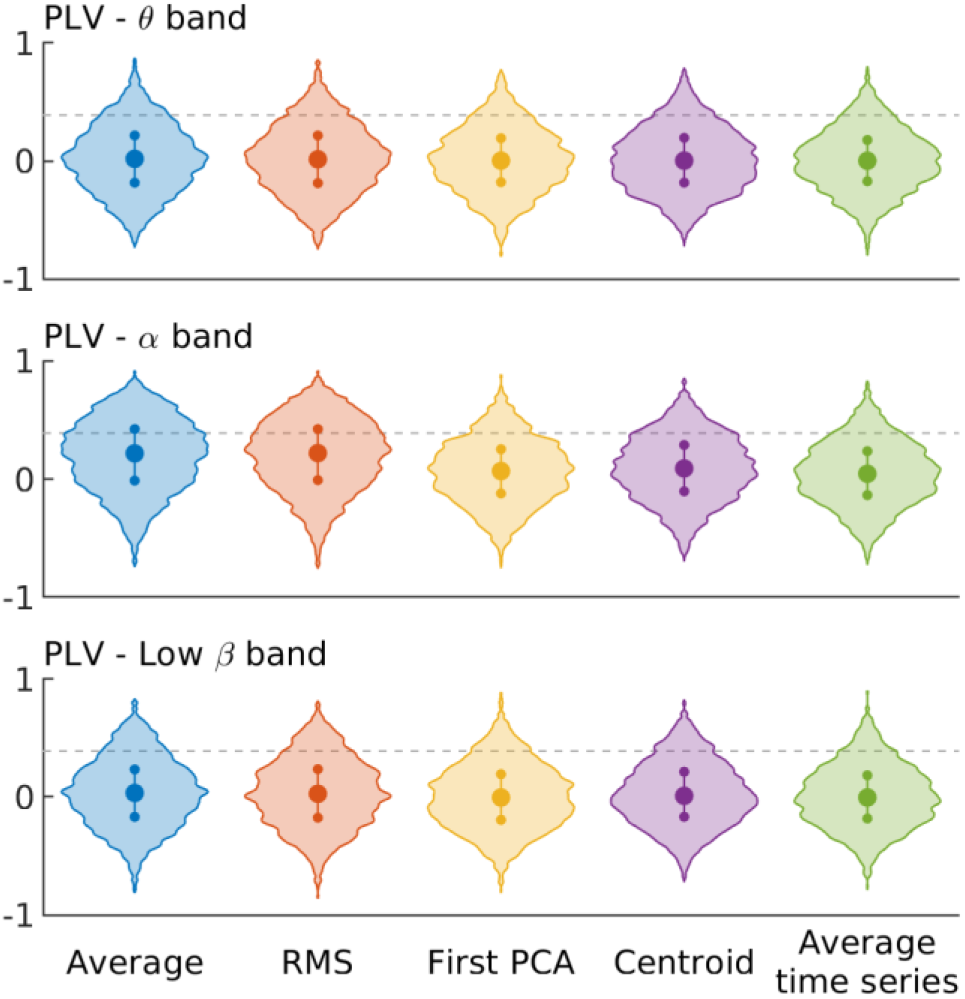
Distribution of correlation coefficients between the PLV estimated using EEG and MEG for each approach. Violin plots showing the correlation between the inter-area PLV calculated with EEG and MEG using different approaches: Blue: Average of the pair-wise PS; Red: Root-mean-square (RMS) of the pair-wise PS; Yellow: PS between the PCA of each area; Purple: PS between the sources closest to the centroid of each area; Green: PS calculated the averaged time series of each area. Each row represents a classical frequency band. Dashed line marks the signification threshold with 5 % (one tailed) alpha level.

### 3.1. PLV and ciPLV using the scalar beamformer

Figure 1 shows the correlation analysis results between EEG and MEG using the classical approach for PLV. Here, each source position is represented by a unique time series, equal to the projection over the direction of maximal activity in the three-dimensional space, achieved using PCA. The results show an almost random distribution of the correlation coefficients for all theta and low beta bands’ approaches and a slightly better than random distribution for the alpha band when using the purely multivariate methods (average and root-mean-square of the pairwise PLV). Table 2 shows the ratio of significant correlation coefficients for each approach and band, supporting this impression.

**Table 2.**
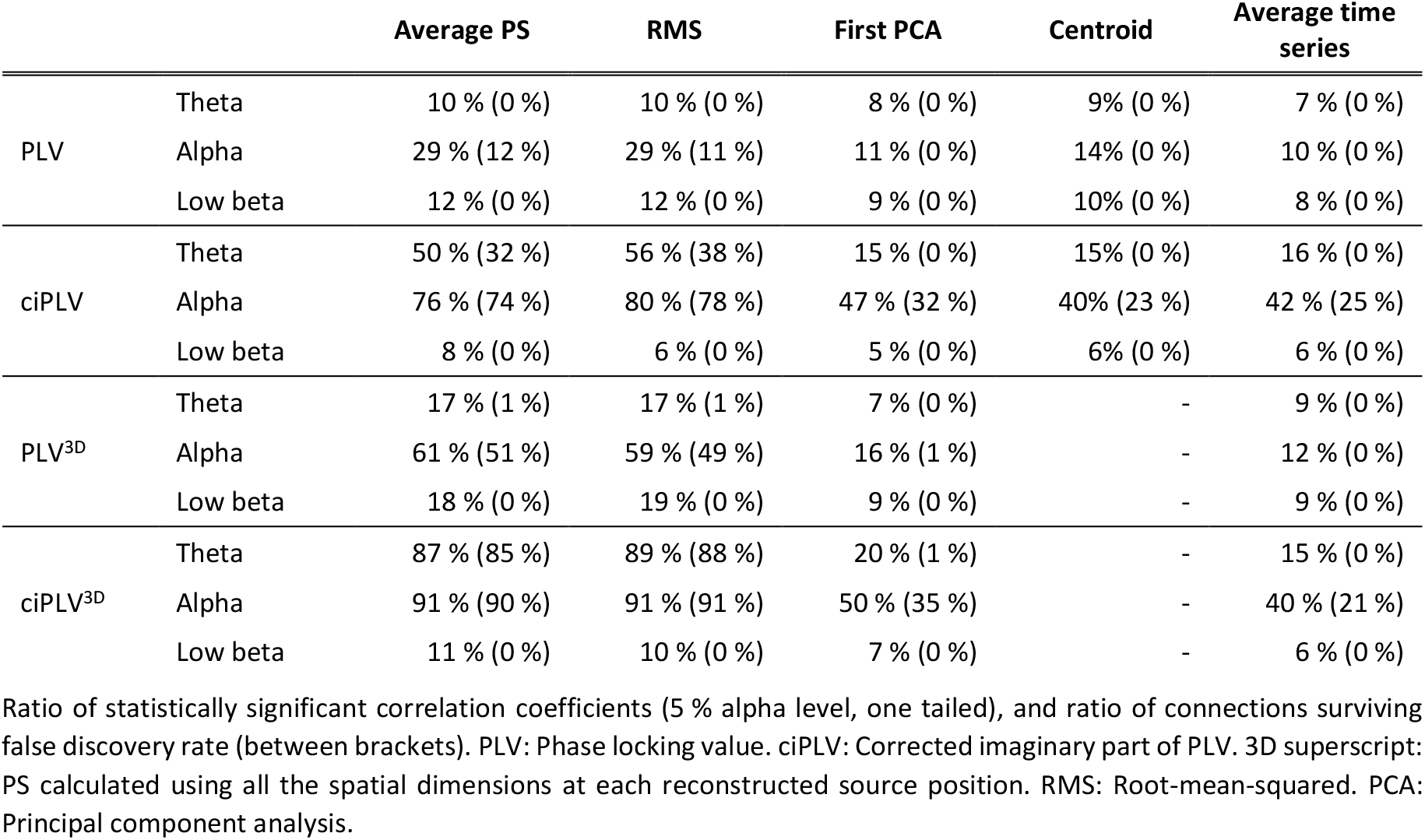
Ratio of statistically significant correlation coefficients between the PS estimated using EEG and MEG for each approach.

Figure 2 shows the same analysis using ciPLV. The low beta band results are still similar to those expected by random data (this can be confirmed by looking at Table 2, where the number of statistically significant correlations is between 5 % and 8 %, and no link survives the false discovery rate). However, the results for the theta and, especially, the alpha band show an interesting behavior. For them, in all cases, the number of statistically significant correlations is above the chance level. The three approaches based on representative time series show similar results, with approximately 15 % (none, after FDR) of statistically significant correlation coefficients in the theta band and around 40 % (20 %, after FDR) in the alpha band. The results for the two purely multivariate approaches are clearly better, with approximately 50 % (30 %, after FDR) and 75 % (70 %, after FDR) of significant correlation coefficients, respectively.

**Figure 2.**
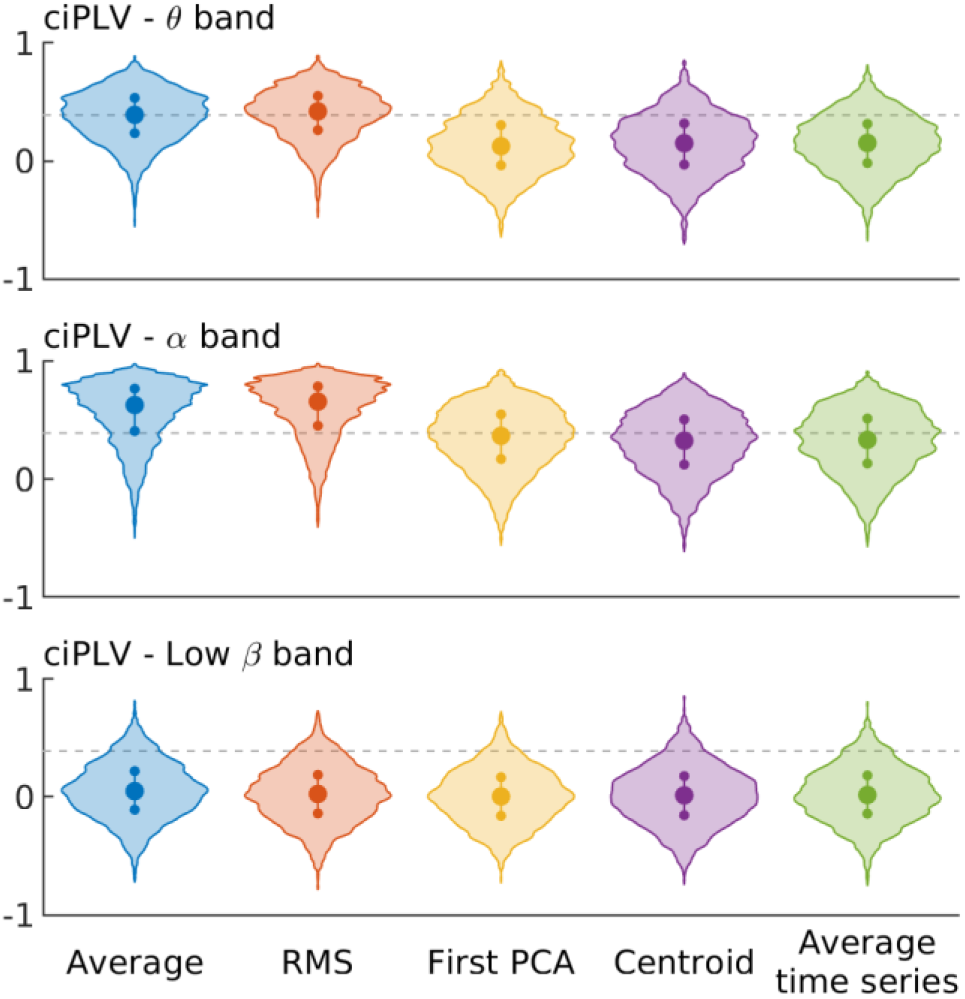
Distribution of correlation coefficients between the ciPLV estimated using EEG and MEG for each approach. Violin plots showing the correlation between the inter-area ciPLV calculated with EEG and MEG using different approaches: Blue: Average of the pair-wise PS; Red: Root-mean-square (RMS) of the pair-wise PS; Yellow: PS between the PCA of each area; Purple: PS between the sources closest to the centroid of each area; Green: PS calculated the averaged time series of each area. Each row represents a classical frequency band. Dashed line marks the signification threshold with 5 % (one tailed) alpha level.

The definition of the significance in a statistical test carries the possibility of false positives, especially when the number of comparisons is high. In order to provide a more understandable view of the correlations between EEG and MEG, we applied a false discovery rate to the results (see 2.6 Statistical analysis). The numerical results are depicted in Table 2, showing a reasonable similitude with the impression described above. When using PLV, all approaches show only random correlations for theta and beta bands, and only the multivariate approaches can identify non-random correlations for the alpha band. On the other hand, when using ciPLV, all the approaches find non-random correlations for the alpha band, but the approaches based on representative time series fail to identify non-random correlations in the theta band.

### 3.2. PLV and ciPLV using the vector (3D) beamformer

However, even the two multivariate approaches discussed above discard some information on brain activity. Indeed, for each source position, we only consider the projection over the orientation of larger activity. Thus, up to 66 % (in the case of EEG) or 50 % (in the case of MEG) of the information could be discarded. To avoid this, we repeated the previous analysis using a different approach, where all the (meaningful) orientations are kept for each source position. The specifics for this computation are described in the Supplementary Materials. As MEG is famously insensitive to one (namely the “radial”) orientation, this approach usually results in three time series per source position in EEG and two in MEG. All the time series belonging to a source position were labeled as belonging to the corresponding area, and then the calculation of the multivariate PS was repeated for each approach. This 3D extension is not possible for the source closest to the centroid of the area (as it will always consist of two or three time series instead of only one “representative” time series), so this approach was not considered.

The results for PLV^3D^ and ciPLV^3D^ are shown in Figure 3. The low beta band results are akin to the ones obtained using only one time series per source position, and again like those expected by chance. The two approaches based on one representative time series per area are similar to those obtained using only one time series per source position in all bands. However, the two purely multivariate approaches show a definite improvement, increasing the ratio of statistically significant correlation coefficients by 70 % (PLV^3D^ in theta band), 100 % (PLV^3D^ in the alpha band), 60 % (ciPLV^3D^ in theta band), and 12 % (ciPLV^3D^ in the alpha band), as shown in Table 2.

**Figure 3.**
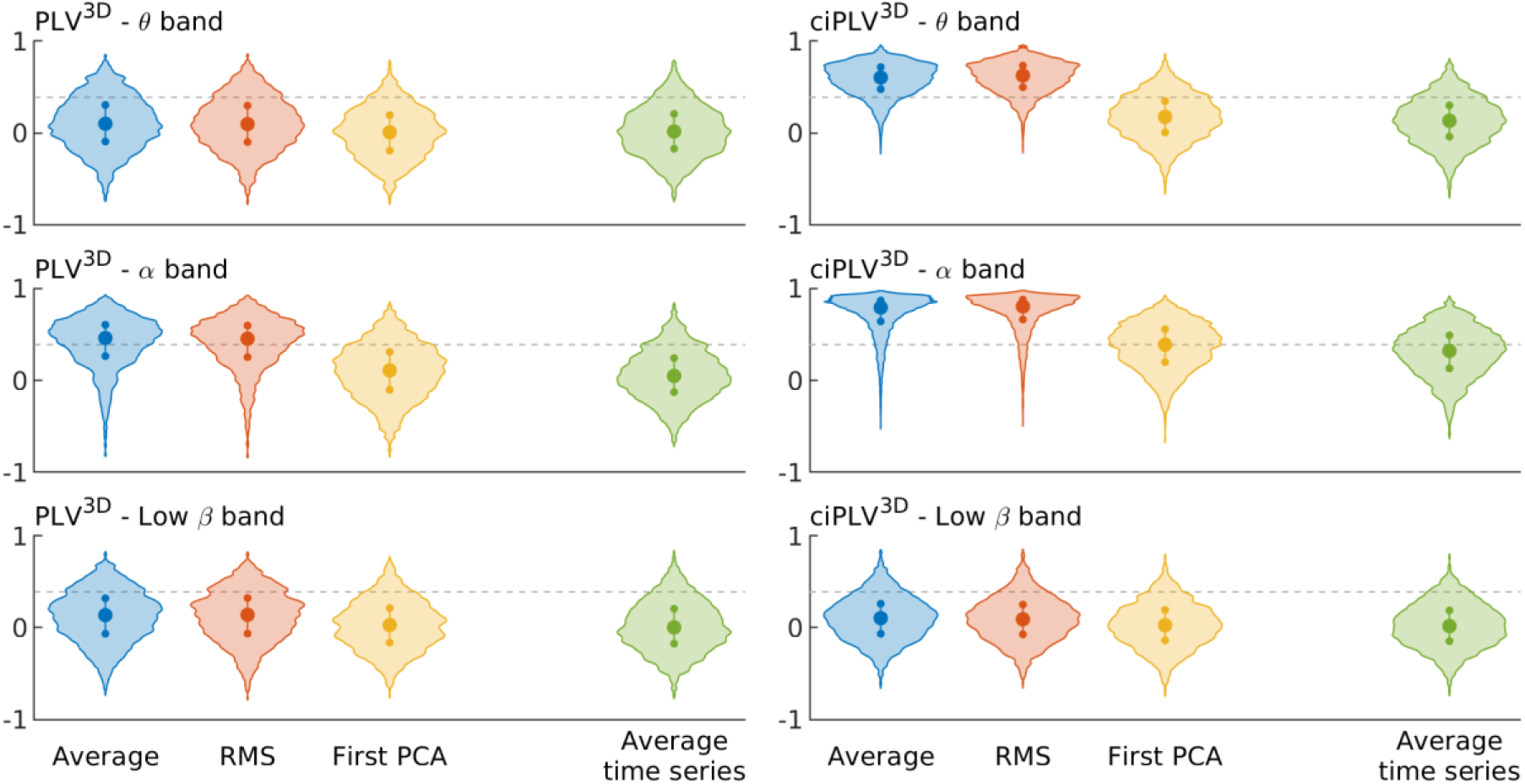
Distribution of correlation coefficients between PLV^3D^ and ciPLV^3D^ estimated using EEG and MEG for each approach. Violin plots showing the correlation between the inter-area PLV^3D^ (left column) and ciPLV^3D^ (right column) calculated with EEG and MEG using all the spatial dimensions and different approaches: Blue: Average of the pair-wise PS; Red: Root-mean-square (RMS) of the pair-wise PS; Yellow: PS between the PCA of each area; Green: PS calculated the averaged time series of each area. Each row represents a classical frequency band. Dashed line marks the signification threshold with 5 % (one tailed) alpha level.

To evaluate the improvement achieved with the 3D versions of PLV and ciPLV, we compared distributions of correlation coefficients using a (one-tailed) related samples *t*-test. The results are depicted, at large, in the Supplementary materials. They show that the improvement is statistically significant for the purely multivariate approaches. In contrast, no such increase is observed for the representative time series based on the area’s average activity. In the case of the representative time series based on the PCA, correlation improves in all cases except for PLV^3D^ in the theta band, where the number of statistically significant correlations actually decreases, as shown in Table 2, from 8 % to 7 % (and in both cases, the number of links surviving FDR is zero).

### 3.3. Hilbert coherence

Hilbert coherence is a variation over PLV and coherence, whose main advantage over PLV is its linear behavior. Therefore, it is interesting to study if the results obtained for PLV and ciPLV are replicable using this new metric. The results are displayed in Figure 4 and are very similar to those obtained using PLV and ciPLV. The most remarkable difference between both metrics can be observed when comparing ciPLV^3D^ and ciHCoh^3D^ in the beta band, as the results for ciHCoh^3D^ show a higher correlation between the FC values estimated using EEG and MEG than those for ciPLV^3D^. The results for all the bands are described, at large, in the Supplementary materials.

**Figure 4.**
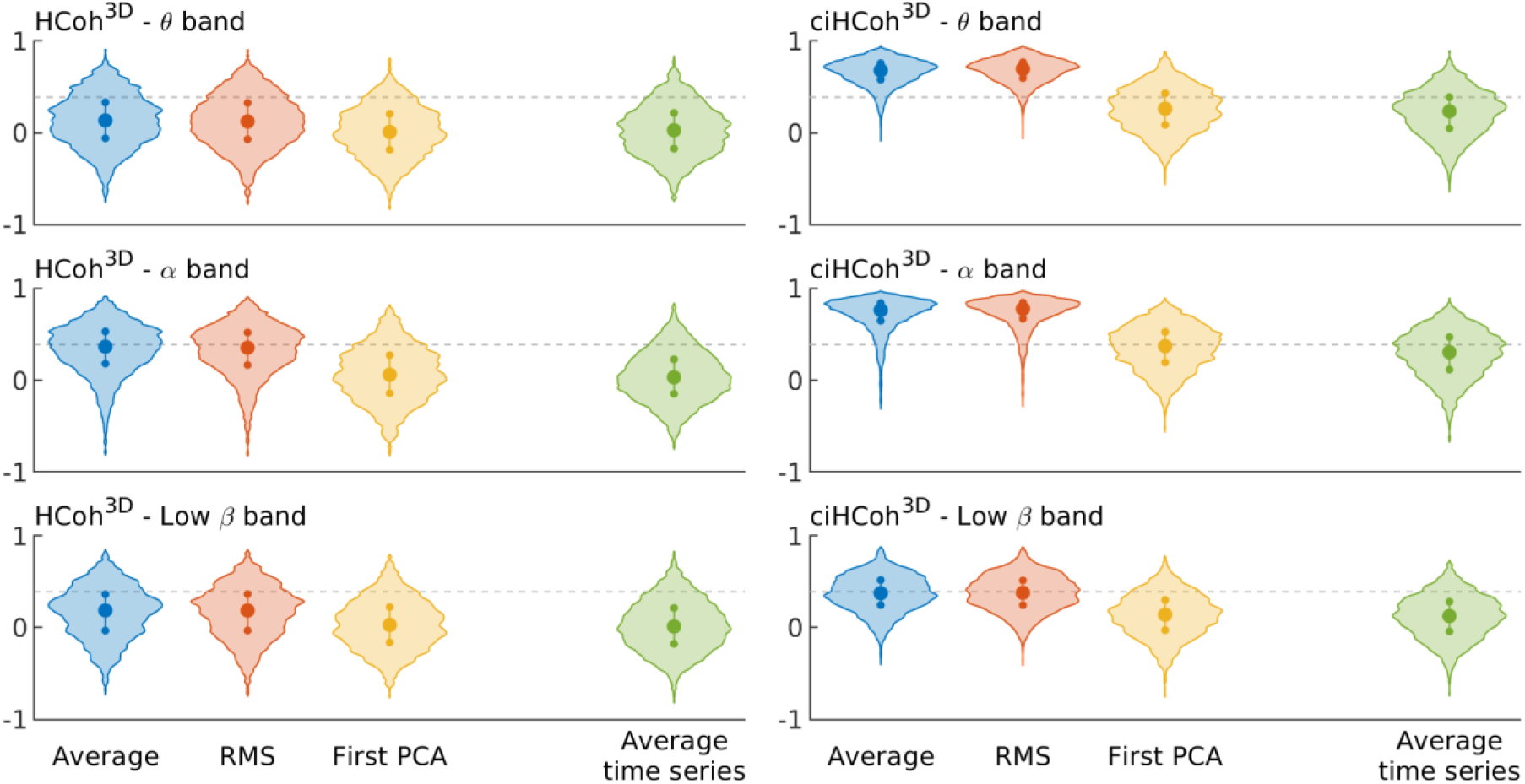
Distribution of correlation coefficients between HCoh^3D^ and ciHCoh^3D^ estimated using EEG and MEG for each approach. Violin plots showing the correlation between the inter-area HCoh^3D^ (left column) and ciHCoh^3D^ (right column) calculated with EEG and MEG using all the spatial dimensions and different approaches: Blue: Average of the pair-wise PS; Red: Root-mean-square (RMS) of the pair-wise PS; Yellow: PS between the PCA of each area; Green: PS calculated the averaged time series of each area. Each row represents a classical frequency band. Dashed line marks the signification threshold with 5 % (one tailed) alpha level.

### 3.4. Spatial distribution of the errors

Finally, to better understand this result, we studied the higher and lower correlations’ spatial distribution. Due to the type of analysis used in this work, it is impossible to identify those areas where the activity is reconstructed with EEG that better match those reconstructed with MEG. Still, we can identify the connectivity links with better matching between both techniques. As it is the approach and band with the best results, we focused this analysis on the alpha band using all the (meaningful) spatial dimensions of the source-space data and the root-mean-square approach. Figure 5 shows the spatial distribution of the 5 % of links with higher (in red) and lower (in blue) correlation between EEG and MEG for PLV^3D^ and ciPLV^3D^. For the PLV^3D^ ones, the links with the highest correlation between EEG and MEG are long projections, while the links with lower correlation are, in general, short ones. For the ciPLV^3D^, on the other hand, the highest correlations appear in posterior areas of the brain, while the lowest ones appear in frontal regions, connecting ventral and orbitofrontal regions.

**Figure 5.**
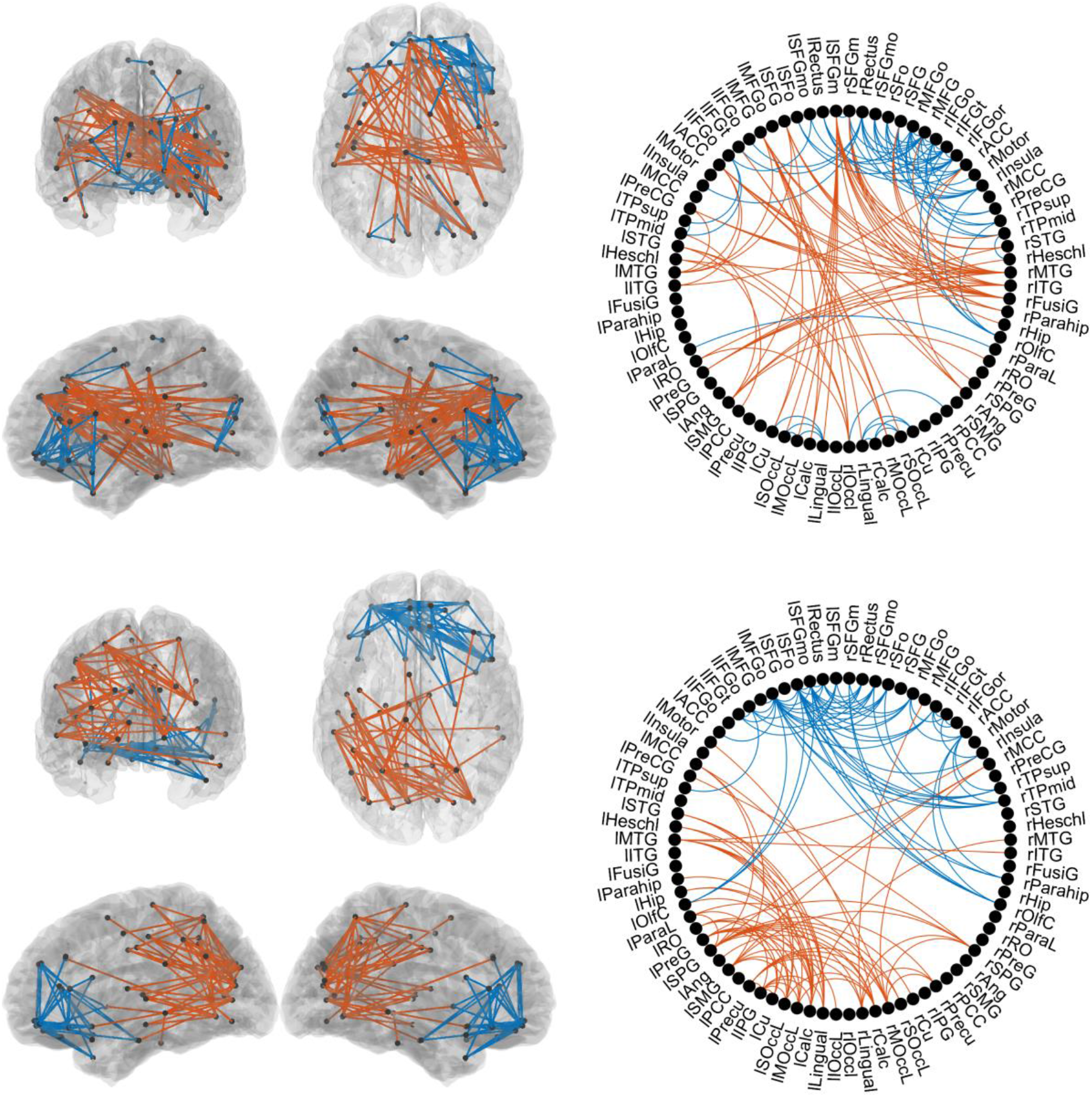
Spatial distribution of the links showing the higher and lower correlation coefficients for PLV^3D^ and ciPLV^3D^ in the alpha band. The figure shows the spatial distribution of FC links with the highest (in red) and the lowest (in blue) correlation between EEG and MEG, when estimated using the RMS approach and PLV^3D^ (upper part) or ciPLV^3D^ (lower part). In both cases, we show the 5 % of links with highest/lowest correlation.

## 4. Discussion

The existence of consistent metrics is mandatory for a correct assessment of functional brain connectivity. Such consistency is crucial when using imperfect electrophysiological recordings such as EEG or MEG because the number of assumptions taken from the data acquisition to the final calculation of source-space synchronization is large. In this work, we have estimated consistency as the similitude in the FC calculated from two different electrophysiological techniques (EEG and MEG) acquired simultaneously and analyzed using the same approaches. For a consistent metric, we expect both results to be similar or, at least, compatible.

Our results clearly show that the purely multivariate approaches (based on the average, or the RMS of the pairwise PS in the links connecting each pair of sources between two areas) are superior to those based on a representative time series of the areas. This result was expected, as the anatomically defined brain areas do not comprise functionally homogeneous sets of sources. Thus, an approach based on taking one time series to represent the whole activity in the area will necessarily discard some information.

Noteworthy, both purely multivariate approaches give similar results, and it is not possible to point out which one shows better behavior. This result is likely due to the limited range of the PS values, in which both means behave similarly. However, from a purely mathematical perspective, the RMS approach is more natural for PLV and ciPLV. PLV is closely related to coherence (Bruña, Maestú, and Pereda, 2018), and coherence is averaged in its squared form (Nunez *et al*., 1997). Aydore and colleagues (Aydore, Pantazis, and Leahy, 2013) showed that many interesting properties of PLV, such as its bias, are closely related to its squared form. Vinck and colleagues also reached the same conclusion when defining the pairwise phase consistency index (Vinck *et al*., 2010). These results suggest that the squared form of PLV shows better mathematical properties than the original form, making sense to average the squared PLV values and then take the square root as a “natural” PLV value. However, this would not necessarily be the case for other PS indexes.

### 4.1. Results for different frequency bands

The correlation between EEG and MEG is, in all cases, higher in the alpha band than in any other band. This outcome could be related to the higher SNR of alpha activity, as it is the dominant brain rhythm when resting with eyes closed. Besides, the alpha band activity seems to consist of one or several oscillators, and the phase synchronization model adapts well to it. This hypothesis is supported by the fact that the low beta band, where no independent oscillator can be (in general) observed, shows the worst performance. In contrast, the high beta band (see the supplementary material), where there is usually a clear oscillator (see, for example, Supplementary material in (Bruña, Maestú, and Pereda, 2018)), gives better correlations.

### 4.2. Effect of zero-lag

When using ciPLV instead of PLV as a metric, the correlations generally improve. This result could be expected, as EEG and MEG show different sensitivities to source leakage. Due to the more significant effect in EEG of imperfections in the head model, EEG source reconstruction is probably more sensitive to source leakage than MEG. When removing the impact of zero-lag synchronization, the effect of source leakage is suppressed (Nolte *et al*., 2004; Stam, Nolte and Daffertshofer, 2007; Brookes, Woolrich and Barnes, 2012; Bruña, Maestú and Pereda, 2018), and the results for EEG and MEG should become more similar. This improvement can be observed in the theta, alpha, high beta, and gamma bands, confirming that low beta does not present coupled oscillatory activity.

However, it is essential to bear in mind that not all zero-lag synchronization is due to source leakage. Real neural dynamics can, in many cases, produce zero-lag interactions, both in the presence of bidirectional communication or through an unseen third element (Kovach, 2017). Besides, the metrics insensitive to zero-lag synchronization have proved, in general, to be less reliable than the ones sensitive to it when evaluated using a test-retest approach (Colclough *et al*., 2016; Garcés, Martín-Buro, and Maestú, 2016). The results in this work should not be considered proof that zero-lag synchronization is spurious, but that the effect of zero lag is *different* in EEG and MEG. When comparing EEG and MEG results, using a metric sensitive to zero-lag synchronization might not be a good idea. However, the information provided by these metrics is still valuable.

### 4.3. Spatial orientations of the source-space activity

The purely multivariate approaches improve when including in the analysis all the (meaningful) orientations of the source space activity instead of using only one projection per source position. If we think of each source position as a small cortical area, the projection over the orientation of maximal activity implies, effectively, taking a representative time series for the source position, and will necessarily entail some sort of information loss. For the case of EEG, when the three-dipole orientations are recorded, up to 66% of the information can be discarded. For MEG, insensitive to the “radial” orientation, the discarded information can be up to 50%. It makes sense that, as happened when considering all pairwise PS values between each pair of sources, including all the possible PS values for the pairwise orientations between each pair of sources improves the estimation of the PS.

### 4.4. Sensitivity to rotations

To allow a correct characterization of the brain dynamics, a good FC estimator must be reliable. Several properties can be studied to quantify a metric’s reliability, and the sensitivity to the coordinate framework is of extreme importance. In the particular case of the brain signal after source reconstruction, this dependency on the coordinate frameworks can be expressed as sensitivity to rotations (Ewald *et al*., 2012; Basti *et al*., 2018). PLV is a nonlinear metric, and it cannot be modified to account for spatial rotations. Namely, PLV cannot be made insensitive to spatial rotations unless the activity is projected over one orientation.

In (3) we defined a linear extension of PLV, the Hilbert coherence. The Supplementary materials show that this new metric’s behavior is closely related to that of the original definition of PLV. Being linear, the Hilbert coherence can be modified (by weighting the contribution of the different orientations; see the Supplementary materials for details) so that it is insensitive to spatial rotations. Thus, this new metric would be insensitive to changes in the coordinate framework. Moreover, this same approach can be applied to the original formula for coherence, resulting in a frequency-wise multivariate estimator for PS, both linear and insensitive to the coordinate framework.

### 4.5. Comparison of EEG and MEG

While not the main objective of this work, the results obtained when comparing EEG and MEG are exciting. MEG is almost blind to radial sources, as the secondary currents create destructive interference in the magnetic field. In the present work, we avoided this issue by projecting the source-space activity over the main directions so that at least 99 % of each source position variance is explained by the resulting time series (see the Supplementary materials for further explanation). This fact implies that, while in EEG, each source’s three components are considered independent, only two components are taken in MEG data. If all the directions were equiprobable, only 66 % of the EEG information could be explained by MEG. However, Figure 3 for the alpha band shows that, at least for non-zero lag, PS, EEG, and MEG share almost the same information.

This result is surprising (Ahlfors *et al*., 2010) but could be explained by the fact that most neurons in the brain are located in the sulci, thereby raising tangential sources. In the Supplementary Materials, we estimate the MEG signal loss due to this insensitivity to radial sources. While these results show that approximately 33 % of the total power of the electrophysiological signal is lost, they do not mean that MEG is blind to one-third of the source positions: MEG is only blind to entirely radial sources, which are a minority of the total. In general, the effect of the insensitivity to the radial orientation of the dipoles is a reduced signal-to-noise ratio. Still, only around 6 % of the cortex is recorded with a sensitivity of 10 % or less (this is, a ten-fold decrease in signal to noise ratio, or more). Instead, our results show that more than 70 % of the cortex is recorded with a sensitivity of 50 % or higher, which increases to 80 % of the cortex if we set the sensitivity limit to 33 %. As the SNR of MEG is better than that of EEG (both because the former is acquired in a low noise environment and has a more accurate and straightforward forward solution), the global SNR of MEG is likely to be similar to that of EEG in most of the cortex. These results imply that, while it is true that MEG is almost blind to radial sources, the effective loss of information derived from this shortcoming is small.

Counterintuitively, this similitude between EEG and MEG does not imply that FC’s estimation from EEG and MEG will give the same, or even similar, results. The FC maps, understood as the relative levels of FC identified between different parts of the brain, estimated from both techniques can, and probably will, differ, as the sensitivity of EEG and MEG for different brain regions is different (see section 4.6 below for an in-depth description of the differences found in this work). Our results indicate that a variation in FC for a given link found in MEG will likely present similarly in EEG, as the FC values estimated for a given connection using EEG and MEG are correlated. For example, if a cognitive task generates an increase in FC between brain region A and brain region B as measured with EEG, it will likely give a similar result when using MEG. This result, of course, depends on the approach used to estimate the FC (and more likely to be true for the multivariate ones), the frequency band (being the correlation stronger in alpha), and the spatial location of the link (being the parietal links more correlated, and the frontal ones less correlated).

### 4.6. Spatial distribution of the differences between EEG and MEG

Figure 5 shows that the links with the highest/lowest correlation between EEG and MEG are not randomly distributed across the brain but clustered. Moreover, its spatial distribution is different for PLV (sensitive to zero-lag synchronization) and ciPLV (insensitive to it).

When using PLV, the links showing the highest EEG/MEG correlation are long connections, including antero-posterior links, inter-hemispheric temporal links, and fronto-temporal links. On the other hand, the lowest correlations appear in short links, especially in frontal and frontal-ventral areas. This result is in all probability related to leakage. Source leakage affects more the estimation of FC between nearby regions, so we would expect the results to show a higher discrepancy between EEG and MEG in this kind of connection when using a metric sensitive to zero-lag synchronization. On the other hand, source leakage is (in general) negligible in connections between distant regions, and we expect these connections to have a similar behavior when measured with EEG and MEG.

The results obtained when using ciPLV are somehow different. The correlation between EEG and MEG seems less affected by the length of the connection, as the effects of source leakage are effectively removed by ignoring the zero-lag synchronization (Stam, Nolte and Daffertshofer, 2007). However, we still find a precise spatial distribution, with the links showing the highest correlation in posterior areas and those showing the lowest correlations (again) in frontal and frontal-ventral areas. The uneven coverage of the head by EEG and MEG can probably explain these results. MEG is notoriously poor at capturing frontal and, especially, fronto-ventral activity, as the sensor array ends in the middle of the forehead (Marinkovic *et al*., 2004). This configuration is required to allow the presentation of visual stimuli to the participants, avoiding covering the eyes with the helmet.

Similarly, EEG’s ventral part of the frontal lobe is poorly covered when using the (extended) 10-20 system (Chatrian, Lettich and Nelson, 1985; Oostenveld and Praamstra, 2001). The most anterior electrodes are the frontal polar ones (Fp1, Fp2, and Fpz), again located in the middle of the forehead, not covering completely the frontal lobe. In this situation, the signal coming from frontal areas should be poorly estimated (both for EEG and MEG), and thus the estimation of FC from these areas would be less reliable than in the rest of the brain.

### 4.7. Computational burden

The main limitation for using the whole pairwise PS matrix is that the computational cost rapidly escalates, compared to the approaches based on representative sources. Using the AAL and focusing only on the 80 cortical brain areas, the number of PLV pairs for these approaches is 3160. On the other hand, using a 1 cm uniform grid, these 80 areas comprise 1210 sources and over 730,000 pairs. When considering all the possible dipolar directions, this number increases to more than 6,580,000 for EEG and 2,925,000 for MEG. However, using the fast PLV approach described in (Bruña, Maestú, and Pereda, 2018), the time required in an average computer to calculate the pairwise PLV matrix (1 kHz sampling rate, 50 trials of 4 seconds, similar to the data used in this work) is less than two minutes per subject and band. Therefore, the whole-brain PS calculation for a cohort of 200 subjects in the five classical bands should take less than a couple of days of computation. The use of Hilbert coherence (a linear extension of PLV) further improves these times, as the calculations take approximately half the time than with PLV, offering similar results. Our group has also derived a set of open-source tools to efficiently estimate PS indices, which can be very useful in this regard (García-Prieto, Bajo, and Pereda, 2017).

### 4.8. Limitations

While we believe our results to be sound, we are aware that our study is not free from some weaknesses. The first one comes from the relatively small number of electrodes in the EEG cap, 32, sometimes considered the bare minimum to perform a decent source reconstruction, as long as an individual MRI is available (Lantz *et al*., 2003; Acar and Makeig, 2013; Sohrabpour *et al*., 2015). On the other hand, after tSSS (Taulu and Simola, 2006), MEG data usually account for approximately 65 independent signals. As we use these sensor-space data to explain 80 cortical regions’ source-space activity, we would expect a lower spatial resolution and, concomitantly, a more significant effect of source leakage in EEG than in MEG (see section 4.6 *Spatial distribution of the differences between EEG and MEG* above). A larger number of electrodes would likely increase spatial resolution and decrease the effects of source leakage.

A second limitation is the use of beamformer as an inverse method. While it is usually considered correct for MEG source reconstruction, it is sometimes disregarded as inadequate for EEG (Gramfort *et al*., 2014). These authors claim that the forward model’s errors, naturally larger in EEG than in MEG, strongly affect beamformer. However, if this were true, the effect we would observe had been a general worsening of the EEG source reconstruction, and the data presented here, especially for ciPLV using all the spatial orientations, indicate otherwise. Besides, even if the source reconstruction in EEG is somehow limited by using beamformers, this limitation would affect all the studied approaches. The goal of this work was to evaluate these different approaches to estimate FC between cortical areas. Therefore, our results indicating which approach is superior are still valid since all of them suffer from a similar penalty due to selecting the inverse method. The evaluation of which inverse method is better for each electrophysiological technique is, on the other hand, beyond the scope of this work. However, it would be interesting, in future works, to study the effect of other extensively used source models (as those based on cortical sheets (Fischl *et al*., 2002)), brain parcellations (for example, the Human Connectome Project atlas (Glasser *et al*., 2016)) and source reconstruction approaches (as the minimum norm (Hämäläinen and Ilmoniemi, 1994) or eLORETA (Pascual-Marqui, 2007a)), in the relation between EEG and MEG.

Lastly, all the participants used in this work are old adults above the age of 60. While there is no reason to think that this fact could alter the results, age-related effects cannot be discarded.

Replication of these analyses in a larger sample covering a broad age range would guarantee our findings’ robustness.

## 5. Conclusions

In this work, we have analyzed the feasibility of calculating PS between brain areas from the activity reconstructed at several source positions distributed across the brain. Our results clearly show that using a multivariate extension of PS, which considers all the possible interactions between the sources of two connected areas, is superior to the (sadly extended) approach of using a unique representative time series per region. Moreover, the estimation quality further improves when using all the spatial orientations of the reconstructed source-space time series, instead of the projection over the direction of maximal activity. We may summarize this in one somewhat tautological but often neglected idea: the more information we use to build a PS estimator, the more accurate the result will be.

The results also show that EEG and MEG are mostly compatible, especially when removing the effects of zero-lag synchronization (and, with them, the impact of source leakage). Even when the FC values estimated are not identical (and we would not expect them to be), the correlation between both techniques is high and well above the statistically significant threshold in many cases. These results indicate that, if a particular brain state, task, or pathology causes an increase (or decrease) in FC, as measured by MEG, between two cortical areas, this same increase (or decrease) will likely appear if using EEG. This outcome is of paramount importance, as it indicates that MEG results are much more transferable to EEG than expected. However, as discussed above, issues such as the sources’ spatial location and the number of channels used in both EEG and MEG must be considered, as their effect on the outcome may be substantial.

Finally, we must note that, as other authors have already pointed out (Marinkovic *et al*., 2004), the quality of the signal, whether EEG or MEG, when measuring frontal and especially frontal-ventral areas, is reduced. Given the importance of frontal cortex in human behavior, some actions should be taken to solve this problem.

## 6. Data and code availability

The data used in this work is available upon request to the correspondence author. Due to privacy concerns, the data cannot be shared freely, and it must be shared in the framework of a collaboration agreement with the Complutense University of Madrid.

The code used in this work is based on FieldTrip (version 20200130) and in-house scripts. The lead fields were built using OpenMEEG 2.2 and in-house scripts. The estimation of the source-space functional connectivity was performed using in-house scripts. All the in-house code used in this work is available at https://github.com/rbruna/the-more-the-merrier.

## Acknowledgments

The authors would like to thank M Gil Poisa for her review of the style of this manuscript. This work was supported by the Ministry of Economy and Competitiveness (MINECO) grant TEC2016-80063-C3-2-R and the Ministry of Science, Innovation, and Universities (MINCIN) grant PID2019-111537GB-C22. RB was partially supported by the Ministry of Education, Culture, and Sport (MECD) grant FPU13/06009.

## References

1. Acar, Z. A. and Makeig, S. (2013) ‘Effects of forward model errors on EEG source localization.’, Brain topography, 26(3), pp. 378–96. doi: 10.1007/s10548-012-0274-6.

2. Ahlfors, S. P. et al. (2010) ‘Sensitivity of MEG and EEG to source orientation.’, Brain topography, 23(3), pp. 227–32. doi: 10.1007/s10548-010-0154-x.

3. Al-Khassaweneh, M. et al. (2016) ‘A Measure of Multivariate Phase Synchrony Using Hyperdimensional Geometry’, IEEE Transactions on Signal Processing, 64(11), pp. 2774–2787. doi: 10.1109/TSP.2016.2529586.

4. Allefeld, C. and Kurths, J. (2004) ‘An approach to multivariate phase synchronization analysis and its application to event-related potentials’, International Journal of Bifurcation and Chaos, 14(02), pp. 417–426. doi: 10.1142/S0218127404009521.

5. Ashburner, J. and Friston, K. J. (2005) ‘Unified segmentation’, NeuroImage, 26(3), pp. 839–851. doi: 10.1016/j.neuroimage.2005.02.018.

6. Aydore, S., Pantazis, D. and Leahy, R. M. (2013) ‘A note on the phase locking value and its properties’, NeuroImage, 74, pp. 231–244. doi: 10.1016/j.neuroimage.2013.02.008.

7. Basti, A. et al. (2018) ‘Disclosing large-scale directed functional connections in MEG with the multivariate phase slope index’, NeuroImage, 175, pp. 161–175. doi: 10.1016/j.neuroimage.2018.03.004.

8. Belouchrani, A. et al. (1997) ‘A blind source separation technique using second-order statistics’, IEEE Transactions on Signal Processing, 45(2), pp. 434–444. doi: 10.1109/78.554307.

9. Benjamini, Y. and Hochberg, Y. (1995) ‘Controlling the false discovery rate: a practical and powerful approach to multiple testing’, Journal of the Royal Statistical Society, 57(1), pp. 289–300. doi: 10.2307/2346101.

10. Biswal, B. et al. (1995) ‘Functional connectivity in the motor cortex of resting human brain using echo-planar MRI’, Magnetic Resonance in Medicine, 34(4), pp. 537–541. doi: 10.1002/mrm.1910340409.

11. Brookes, M. J., Woolrich, M. W. and Barnes, G. R. (2012) ‘Measuring functional connectivity in MEG: A multivariate approach insensitive to linear source leakage’, NeuroImage, 63(2), pp. 910–920. doi: 10.1016/j.neuroimage.2012.03.048.

12. Bruña, R., Maestú, F. and Pereda, E. (2018) ‘Phase locking value revisited: Teaching new tricks to an old dog’, Journal of neural engineering. Institute of Physics Publishing, 15(5), p. 056011. doi: 10.1088/1741-2552/aacfe4.

13. Chatrian, G. E., Lettich, E. and Nelson, P. L. (1985) ‘Ten percent electrode system for topographic studies of spontaneous and evoked EEG activity’, American Journal Of EEG Technology, 25(2), pp. 83–92. doi: 10.1080/00029238.1985.11080163.

14. Colclough, G. L. et al. (2016) ‘How reliable are MEG resting-state connectivity metrics?’, NeuroImage, 138, pp. 284–293. doi: 10.1016/j.neuroimage.2016.05.070.

15. Craddock, R. C. et al. (2012) ‘A whole brain fMRI atlas generated via spatially constrained spectral clustering’, Human Brain Mapping, 33(8), pp. 1914–1928. doi: 10.1002/hbm.21333.

16. Desikan, R. S. et al. (2006) ‘An automated labeling system for subdividing the human cerebral cortex on MRI scans into gyral based regions of interest’, NeuroImage, 31, pp. 968–980. doi: 10.1016/j.neuroimage.2006.01.021.

17. Ewald, A. et al. (2012) ‘Estimating true brain connectivity from EEG/MEG data invariant to linear and static transformations in sensor space’, NeuroImage, 60(1), pp. 476–488. doi: 10.1016/j.neuroimage.2011.11.084.

18. Fang, Q. and Boas, D. A. (2009) ‘Tetrahedral mesh generation from volumetric binary and grayscale images’, in Proceedings - 2009 IEEE International Symposium on Biomedical Imaging: From Nano to Macro, ISBI 2009, pp. 1142–1145. doi: 10.1109/ISBI.2009.5193259.

19. Fischl, B. R. et al. (2002) ‘Whole brain segmentation: automated labeling of neuroanatomical structures in the human brain.’, Neuron, 33(3), pp. 341–55.

20. Friston, K. J. (1994) ‘Functional and effective connectivity in neuroimaging: A synthesis’, Human Brain Mapping, 2(1–2), pp. 56–78. doi: 10.1002/hbm.460020107.

21. Friston, K. J. et al. (2006) ‘A critique of functional localisers.’, NeuroImage, 30(4), pp. 1077–1087. doi: 10.1016/j.neuroimage.2005.08.012.

22. Garcés, P. et al. (2017) ‘Choice of magnetometers and gradiometers after signal space separation’, Sensors (Switzerland), 17(12), p. 2926. doi: 10.3390/s17122926.

23. Garcés, P., Martín-Buro, M. C. and Maestú, F. (2016) ‘Quantifying the Test-Retest Reliability of Magnetoencephalography Resting-State Functional Connectivity’, Brain Connectivity, 6(6), pp. 448–460. doi: 10.1089/brain.2015.0416.

24. García-Prieto, J., Bajo, R. and Pereda, E. (2017) ‘Efficient Computation of Functional Brain Networks: toward Real-Time Functional Connectivity.’, Frontiers in neuroinformatics, 11, p. 8. doi: 10.3389/fninf.2017.00008.

25. Geschwind, N. (1965) ‘Disconnexion syndromes in animals and man. Part I’, Brain, 88(2), pp. 237–294. doi: 10.1007/s11065-010-9131-0.

26. Glasser, M. F. et al. (2016) ‘A multi-modal parcellation of human cerebral cortex.’, Nature, 536(7615), pp. 171–178. doi: 10.1038/nature18933.

27. Gramfort, A. et al. (2010) ‘OpenMEEG: opensource software for quasistatic bioelectromagnetics.’, Biomedical engineering online, 9, p. 45. doi: 10.1186/1475-925X-9-45.

28. Gramfort, A. et al. (2014) ‘MNE software for processing MEG and EEG data’, NeuroImage, 86, pp. 446–460. doi: 10.1016/j.neuroimage.2013.10.027.

29. Gross, J. et al. (2001) ‘Dynamic imaging of coherent sources: Studying neural interactions in the human brain.’, Proceedings of the National Academy of Sciences of the United States of America, 98, pp. 694–699. doi: 10.1073/pnas.98.2.694.

30. Groth, A. and Ghil, M. (2011) ‘Multivariate singular spectrum analysis and the road to phase synchronization’, Physical Review E - Statistical, Nonlinear, and Soft Matter Physics, 84, p. 036206. doi: 10.1103/PhysRevE.84.036206.

31. Hämäläinen, M. S. and Ilmoniemi, R. J. (1994) ‘Interpreting magnetic fields of the brain: minimum norm estimates’, Medical & Biological Engineering & Computing, 32(1), pp. 35–42. doi: 10.1007/BF02512476.

32. Hincapié, A.-S. et al. (2017) ‘The impact of MEG source reconstruction method on source-space connectivity estimation: A comparison between minimum-norm solution and beamforming’, NeuroImage, 156, pp. 29–42. doi: 10.1016/j.neuroimage.2017.04.038.

33. Hotelling, H. (1936) ‘Relations between two sets of variates’, Biometrika, 28(3–4), pp. 321–377. doi: 10.1093/biomet/28.3-4.321.

34. James, G. A., Hazaroglu, O. and Bush, K. A. (2016) ‘A human brain atlas derived via n-cut parcellation of resting-state and task-based fMRI data’, Magnetic Resonance Imaging, 34(2), pp. 209–218. doi: 10.1016/j.mri.2015.10.036.

35. King, D. R. et al. (2015) ‘Recollection-Related Increases in Functional Connectivity Predict Individual Differences in Memory Accuracy’, Journal of Neuroscience, 35(4), pp. 1763–1772. doi: 10.1523/JNEUROSCI.3219-14.2015.

36. Korhonen, O., Palva, S. and Palva, J. M. (2014) ‘Sparse weightings for collapsing inverse solutions to cortical parcellations optimize M/EEG source reconstruction accuracy’, Journal of Neuroscience Methods, 226, pp. 147– 160. doi: 10.1016/j.jneumeth.2014.01.031.

37. Kovach, C. K. (2017) ‘A Biased Look at Phase Locking: Brief Critical Review and Proposed Remedy’, IEEE Transactions on Signal Processing, 65(17), pp. 4468–4480. doi: 10.1109/TSP.2017.2711517.

38. Kwong, K. K. et al. (1992) ‘Dynamic magnetic resonance imaging of human brain activity during primary sensory stimulation.’, Proceedings of the National Academy of Sciences of the United States of America, 89(12), pp. 5675–5679. doi: 10.1073/pnas.89.12.5675.

39. Lachaux, J.-P. et al. (1999) ‘Measuring phase synchrony in brain signals.’, Human brain mapping, 8, pp. 194– 208. doi: 10.1002/(SICI)1097-0193(1999)8:4<194::AID-HBM4>3.0.CO;2-C.

40. Lantz, G. et al. (2003) ‘Epileptic source localization with high density EEG: how many electrodes are needed?’, Clinical Neurophysiology, 114(1), pp. 63–69. doi: 10.1016/S1388-2457(02)00337-1.

41. Lohmann, G. et al. (2010) ‘Eigenvector centrality mapping for analyzing connectivity patterns in fMRI data of the human brain’, PLoS ONE, 5(4), p. e10232. doi: 10.1371/journal.pone.0010232.

42. López-Sanz, D. et al. (2017) ‘Network Disruption in the Preclinical Stages of Alzheimer’s Disease: From Subjective Cognitive Decline to Mild Cognitive Impairment’, International Journal of Neural Systems, 27(08), p. 1750041. doi: 10.1142/S0129065717500411.

43. Luckhoo, H. et al. (2012) ‘Inferring task-related networks using independent component analysis in magnetoencephalography’, NeuroImage, 62(1), pp. 530–541. doi: 10.1016/j.neuroimage.2012.04.046.

44. Marinkovic, K. et al. (2004) ‘Head position in the MEG helmet affects the sensitivity to anterior sources.’, Neurology & clinical neurophysiology : NCN, 2004, p. 30. Available at: http://www.ncbi.nlm.nih.gov/pubmed/16012659.

45. Mesulam, M. M. (2015) ‘Fifty years of disconnexion syndromes and the Geschwind legacy’, Brain, 138(9), pp. 2791–9. doi: 10.1093/brain/awv198.

46. Mormann, F. et al. (2000) ‘Mean phase coherence as a measure for phase synchronization and its application to the EEG of epilepsy patients’, Physica D: Nonlinear Phenomena, 144(3), pp. 358–369. doi: 10.1016/S0167-2789(00)00087-7.

47. Nolte, G. et al. (2004) ‘Identifying true brain interaction from EEG data using the imaginary part of coherency’, Clinical Neurophysiology, 115(10), pp. 2292–2307. doi: 10.1016/j.clinph.2004.04.029.

48. Nolte, G. et al. (2008) ‘Robustly estimating the flow direction of information in complex physical systems’, Physical Review Letters, 100(23), p. 234101. doi: 10.1103/PhysRevLett.100.234101.

49. Nunez, P. L. et al. (1997) ‘EEG coherency I: Statistics, reference electrode, volume conduction, Laplacians, cortical imaging, and interpretation at multiple scales’, Electroencephalography and Clinical Neurophysiology, 103(5), pp. 499–515. doi: 10.1016/S0013-4694(97)00066-7.

50. Oostenveld, R. et al. (2011) ‘FieldTrip: Open Source Software for Advanced Analysis of MEG, EEG, and Invasive Electrophysiological Data’, Computational Intelligence and Neuroscience, 2011, pp. 1–9. doi: 10.1155/2011/156869.

51. Oostenveld, R. and Praamstra, P. (2001) ‘The five percent electrode system for high-resolution EEG and ERP measurements’, Clinical Neurophysiology, 112(4), pp. 713–719. doi: 10.1016/S1388-2457(00)00527-7.

52. Pascual-Marqui, R. D. (2007a) ‘Discrete, 3D distributed, linear imaging methods of electric neuronal activity. Part 1: exact, zero error localization’, arXiv. Available at: http://arxiv.org/abs/0710.3341.

53. Pascual-Marqui, R. D. (2007b) ‘Instantaneous and lagged measurements of linear and nonlinear dependence between groups of multivariate time series: frequency decomposition’, arXiv. Available at: http://arxiv.org/abs/0711.1455.

54. Rosenblum, Pikovsky and Kurths (1996) ‘Phase synchronization of chaotic oscillators.’, Physical review letters, 76(11), pp. 1804–1807. doi: 10.1103/PhysRevLett.76.1804.

55. Schwabedal, J. T. C. and Kantz, H. (2016) ‘Optimal Extraction of Collective Oscillations from Unreliable Measurements.’, Physical review letters, 116(10), p. 104101. doi: 10.1103/PhysRevLett.116.104101.

56. Sohrabpour, A. et al. (2015) ‘Effect of EEG electrode number on epileptic source localization in pediatric patients’, Clinical Neurophysiology, 126(3), pp. 472–480. doi: 10.1016/j.clinph.2014.05.038.

57. Sporns, O., Tononi, G. and Kötter, R. (2005) ‘The human connectome: A structural description of the human brain.’, PLoS Computational Biology, 1(4), p. e42. doi: 10.1371/journal.pcbi.0010042.

58. Stam, C. J., Nolte, G. and Daffertshofer, A. (2007) ‘Phase lag index: Assessment of functional connectivity from multi channel EEG and MEG with diminished bias from common sources’, Human Brain Mapping, 28(11), pp. 1178–1193. doi: 10.1002/hbm.20346.

59. Taulu, S. and Simola, J. (2006) ‘Spatiotemporal signal space separation method for rejecting nearby interference in MEG measurements’, Physics in Medicine and Biology, 51(7), pp. 1759–1768. doi: 10.1088/0031-9155/51/7/008.

60. Tzourio-Mazoyer, N. et al. (2002) ‘Automated anatomical labeling of activations in SPM using a macroscopic anatomical parcellation of the MNI MRI single-subject brain.’, NeuroImage, 15(1), pp. 273–89. doi: 10.1006/nimg.2001.0978.

61. van Veen, B. D. et al. (1997) ‘Localization of brain electrical activity via linearly constrained minimum variance spatial filtering.’, IEEE transactions on bio-medical engineering, 44, pp. 867–880. doi: 10.1109/10.623056.

62. Vinck, M. et al. (2010) ‘The pairwise phase consistency: a bias-free measure of rhythmic neuronal synchronization.’, NeuroImage, 51(1), pp. 112–22. doi: 10.1016/j.neuroimage.2010.01.073.

