## Supplementary Materials for "The more, the merrier: multivariate phase synchronization methods excel pairwise ones in estimating functional brain connectivity from reconstructed neural sources"

*Calculation of the meaningful orientations at each source position*

This work's source model comprised 1210 cortical sources in a homogeneous grid built in the MNI template, with 1 cm of spacing. As there are no anatomical constraints to each source position's orientation, the three spatial dimensions are studied using three orthogonal dipoles per source position. For the first part of the study, a unique representative time series per source position was considered, representing the source activity's spatial projection over the direction of maximal activity. It was calculated as the main component, obtained using principal component analysis (PCA), of the three possible orientations. This approach has the downside of discarding up to 50 % (in MEG) or 66 % (in EEG) of the activity, as the second and (in its case) third components are not included.

As our goal was to include all the information at each source position in the calculation of the inter-area PS, we repeated the analysis using all the meaningful orientations at each source position. This analysis is referred to as PLV 3D, but it does not necessarily include all three orientations. As shown above, in MEG, each source position is defined only by two orthogonal orientations, as this technique is blind to the "radially" oriented dipoles. However, due to numerical inaccuracies, the reconstructed time series has three dimensions, one of them negligible in terms of power compared to the other two. This is usually not an issue, as the contribution of this "noisy" component is low. However, the calculation of PLV uses only the signal phase, discarding its amplitude, so the contributions of all three orientations are weighted equally, artificially increasing the effect of noise.

We avoided this issue by discarding the "radial" component at each source position for MEG. This aim was achieved systematically using PCA: the activity at each source position was projected over the (orthogonal) directions of the higher, second to higher, and lower activity, this last component corresponding to the radial orientation. Then it was discarded, using as a criterion that the remaining components should comprise 99 % of the total variance. This approach guarantees that all the meaningful information (up to that 99 % of activity) is kept while removing the noise. The same procedure was used for EEG, guaranteeing that no noisy components are included in the PS's computation. After this projection, the resulting source-space data consisted of two orientations per source position in MEG and between one and three orientations per source position (three in more than 97 % of the cases) for EEG.

*MEG sensitivity to source orientation*

Magnetoencephalography is famously known to be blind to some source orientations. When using a spherical head model, this technique is blind to the radial component of the primary current (Sarvas, 1987), as the magnetic field generated by the symmetrically propagating secondary currents cancels out the one produced by the primary current. One could think that this would not be the case for more realistic head models, but Ahlfors and colleagues proved that, for every source position in any head model, there is a dipole orientation to which MEG is blind (Ahlfors *et al.*, 2010). This orientation is usually referred to as the radial orientation, even in realistic head models. EEG does not present this weakness and is equally sensitive to every dipolar orientation.

For a brain activity consisting of a distribution of sources with random orientations, MEG's insensitivity to radial sources would imply that the agreement between MEG and EEG could never be above 66% of the signal variance. However, the generators of electrophysiological activity in the brain do not obey a random distribution. Simulations show that most of the signals captured by these techniques are generated by the presynaptic currents at the pyramidal neurons of the cortex (Murakami and Okada, 2006). The apical dendrite of these neurons, leading contributors to these currents, lies perpendicular to the cortical surface. Due to the cerebral cortex's intricate structure, most of the neurons are located in the sulci walls, with only a small fraction of them located in the gyri or at the sulci's bottom. Most of the cortex's neurons present orientations *tangential* to the inner skull interface and may be considered tangential dipoles.

Supplementary Figure 1. Sensitivity of the MEG across the cortex for participant mq-01.

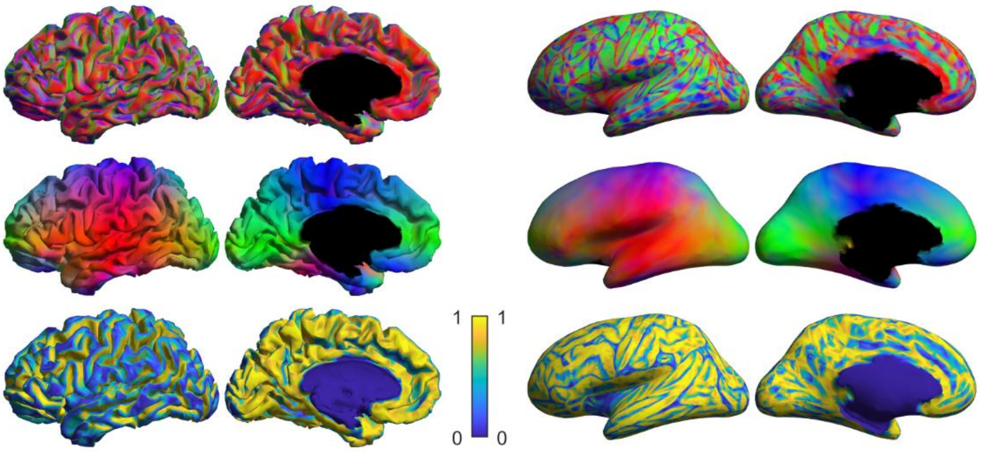


The figure shows the sensitivity of the MEG to each region of the cortex. The top row shows the theoretical orientation of the electrophysiological current, normal to the surface of the cortex; red color indicates orientation with the X (left/right) axis, green color indicates orientation with the Y (front/back) axis, and blue color indicates orientation with the Z (up/down) axis. The middle row shows the blind orientation (the "radial" orientation) at each point of the cortex; the color scheme is the same than in the top row. The bottom row shows the sensitivity of the MEG to each region of the cortex, in per-one units of power. Left part of the figure shows the original cortical surface, and right part shows the inflated surface. Please, note that neither representation is accurate in terms of sensitivity per unit of area: the direct representation hides the sulci, and the inflated representation warps the surface.

To get a better quantification of the radial sources, we obtained the cortical surface, using FreeSurfer (Fischl *et al.*, 2002), for 8 of our participants who had an available 3T T1 MRI image. We calculated the normal of this cortical surface at each node to estimate the primary currents and selected approximately 15,000 evenly distributed cortical points using a mesh simplification. Then, for each source position, we calculated the "radial orientation" using a PCA decomposition of the MEG lead field and estimated the projection of the theoretical current (perpendicular to the cortex) over this orientation. This procedure allows us to determine the loss of sensitivity due to this "blindness" of MEG. The results of this analysis for one single participant (mq-01) are shown in Supplementary Figure 1. The figure shows the expected sensitivity map, with high MEG sensitivity in the sulci and low sensitivity in the gyri. However, the interpretation of the ratio of sources with high and low sensitivity is not straightforward, as the cortex's complex structure creates distortion when representing the inflated brain surface.

A more accurate estimation of the overall MEG sensitivity would include the cortex's sensitivity per unit area. In squared millimeters, we calculated the cortical area, represented for each of the (approximately) 15,000 source positions, and assigned to this area the central source's sensitivity. This procedure results in a map showing the sensitivity per unit area. If we consider that the source density is homogeneous across the cortex, the result shows the MEG system's overall sensitivity to the source activity. The results are shown in Supplementary Figure 2, in the form of violin-plots representing the brain's sensitivity distribution per unit area. They demonstrate that, overall, approximately 33 % of the signal power is lost due to MEG's blindness to radial sources. Still, this loss is distributed and not concentrated in a small number of sources. That is, MEG is not blind to one-third of the sources in the brain, but to one-third of the total power. The result of the insensitivity to radial sources is a general loss of signal-to-noise ratio. Moreover, this loss is not homogeneous, and more than 75 % of brain sources are acquired with a higher than 45 % sensitivity. In contrast, only 6 % of the sources are acquired with sensitivity lower than 10 %.

Supplementary Figure 2. Distribution of sensitivity per unit of area for each participant.

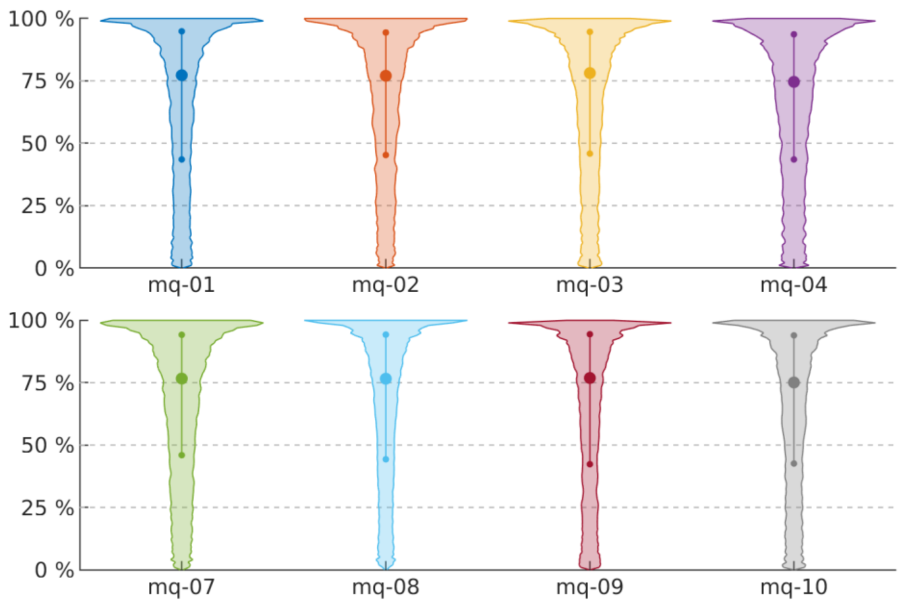


Violin plots showing the distribution of sensitivity (in per-cent units of source power) per unit of cortical area for the eight participants with a 3D T1 MRI image. Dashed lines represent the points of 75 %, 50 %, and 25 % sensitivity. The dots inside the violins represent the first, second (median) and third percentile of the distribution. Approximately 50 % of the cortical area, for each individual, is recorded with a sensitivity of 75 % or higher, and approximately 75 % of the cortical area is recorded with a sensitivity of 40 % or higher.

*Sensitivity to rotations*

To allow for a correct characterization of the brain dynamics, a good FC estimator must be reliable. Several properties can be studied to quantify a metric's reliability, and the sensitivity to the coordinate frame is one of extreme importance. In the brain signal's particular case after source reconstruction, this dependency on the coordinate system can be expressed as sensitivity to rotations (Ewald *et al.*, 2012; Basti *et al.*, 2018). The brain generators of the EEG and MEG signal in a given point of the brain are multidimensional, with two (for MEG, see *MEG orientations* above) or three (in the case of EEG) perpendicular components with arbitrary orientations with respect to the reference frame. If we aim to estimate the real FC between two points in the brain, all these dimensions must somehow be taken into account.

In the case of source reconstruction methods where the output per source location is scalar, the source model with orientation constraints (Dale and Sereno, 1993) or SAM beamformer (Vrba and Robinson, 2001), the method itself is insensitive to rotations. Similarly, when the vector-form source reconstruction is projected over the direction of maximal activity, the result is also insensitive to rotations of the coordinate frame. However, it could be argued that, as the real nature of the signal generators is not unidimensional, these methods imply a loss of information. Indeed, the results of this work seem to confirm this intuition.

In this work, we calculated the brain activity using all the meaningful orientations (see above) defined by PCA. With this procedure, we define a new coordinate framework dependent on the data so that the first direction is that of maximal activity, the third direction is that of minimal activity, and the second direction is perpendicular to both. As each source position's activity is rotated using this criterion, the original coordinate system is irrelevant, and an eventual rotation on it will be compensated. However, in a more general scenario, where the source model arbitrarily defines the three orientations at each source, they must be integrated consistently. In the case of Hilbert coherence, this can be performed trivially. Hilbert coherence can be interpreted as (the absolute value of) the correlation of the Hilbert analytical signals, and the correlation corresponds to the normalized covariance. Thus, Hilbert coherence can be written in terms of the covariance:

$$HCoh = \mathrm{corr}\big(\tilde{x}(t), \tilde{y}(t)\big) = \frac{\mathrm{cov}\big(\tilde{x}(t), \tilde{y}(t)\big)}{\mathrm{var}\big(\tilde{x}(t)\big) \cdot \mathrm{var}\big(\tilde{y}(t)\big)} \quad (1)$$

When working with three-dimensional time series, the covariance is represented as a 3-by-3 matrix, and rotation in these time series implies rotation over this matrix:

$$\mathrm{cov}\big(r_1 \cdot s_1(t), r_2 \cdot s_2(t)\big) = r_1 \cdot \mathrm{cov}\big(s_1(t), s_2(t)\big) \cdot r_2^* \quad (2)$$

where $s_\mathrm{i}(t)$ is the $i$-*th* three-dimensional time series, and $r_\mathrm{i}$ the corresponding 3-by-3 rotation matrix. As the Frobenius norm is invariant to rotations, the original covariance matrix norm is equal to the rotated one. This norm can be interpreted as the covariance of two three-dimensional time series, which must be normalized to get the corresponding correlation coefficient. A meaningful normalization factor would be the product of the total power at each source position. In the case of a one-dimension time series, the total power equals the time series variance itself, and the result equals (1). In the case of a multidimensional time series, the total power equals the trace of the covariance matrix:

$$wHCoh = \frac{\left\| \mathrm{cov}\left(\widetilde{s_1}(t), \widetilde{s_2}(t)\right) \right\|}{\mathrm{tr}\left\{\mathrm{cov}\left(\widetilde{s_1}(t)\right)\right\} \cdot \mathrm{tr}\left\{\mathrm{cov}\left(\widetilde{s_2}(t)\right)\right\}} \tag{3}$$

where $\|\cdot\|$ and $\mathrm{tr}\{\cdot\}$ stand for the Frobenius norm and the trace, respectively. As the trace of a covariance matrix is invariant to rotations, the correlation calculated in this way would also be invariant to rotations. We can name this metric weighted HCoh (wHCoh) because the result is equivalent to a weighted RMS of the elements in HCoh, being the weights equal to the power corresponding to each entry in the matrix. wHCoh is invariant to rotations, giving a more meaningful result in the general case. However, it requires two calculations: first, the weighted RMS of the HCoh matrices for each pair of source positions, and second the per-area average. As our methodology, based on the projection of the time series using PCA, is already insensitive to rotations, we have not used this metric in the current work.

**On the term "multivariate phase synchronization."**

Once the sources of neural activity have been distributed into the different brain areas, estimating the degree of FC between any two of these areas (say, $A$ and $B$) consists of mapping the $N_A \times N_B$ time series of length M to a single real number, $\rho_{AB}$, between 0 and 1. Namely, the FC estimator is a function, $f$, acting on the space of as follows:

$$\mathbb{R}^{N_A \times M} \times \mathbb{R}^{N_B \times M} \xrightarrow{f} [0,1] \tag{4}$$

where $\mathbb{R}^{N_A \times M}$ (resp. $\mathbb{R}^{N_B \times M}$) is the space of real-valued matrices of $\boldsymbol{A}, \boldsymbol{B}$ of dimension $N_A \times M$ (areas x samples). This mapping entails a considerable reduction of dimensionality yet defining the best $f(A, B)$ is far from trivial.

*Bivariate methods*

In the case where we decide to choose (or construct) a single representative time series for each area, FC estimation takes the following form:

$$\mathbb{R}^{N_A \times M} \times \mathbb{R}^{N_B \times M} \xrightarrow{g} \mathbb{R}^{1 \times M} \times \mathbb{R}^{1 \times M} \xrightarrow{h} [0,1] \tag{5}$$

Here, function $g$ performs the initial reduction transforming matrices $\boldsymbol{A}$ and $\boldsymbol{B}$ into two real-valued time series (vectors) $A$ and $B$ of dimension $1 \times M$, whereas $h$ is the function that estimates the FC index. The function $f$ is therefore defined here as $f = h\big(g(\boldsymbol{A}, \boldsymbol{B})\big) = h(A, B)$. In other words, $f$ represents a two-stage procedure: the initial reduction of the dimensionality (in the time domain, function $g$) produces one time series per area. Moreover, the FC between the areas is estimated using a bivariate index (function $h$) operating on these resulting univariate time series.

Note that the function $g$ that produces the representative time series can take many forms. In the simplest, most frequently case where it only considers the sources in the corresponding area but not those on the other one, we have $\mathbb{R}^{N_Y \times M} \xrightarrow{g} \mathbb{R}^{1 \times M}$, where $Y$ stands for all the atlas areas. Usual choices for $g$ are the average or the PCA of all the sources or may imply choosing a single source such as the centroid or the one with the greatest power. A more sophisticated version of $g$ in (5) uses the information of the sources in both areas. For instance, in the classic canonical correlation analysis (CCA), $g$ generates the vectors $A$ and $B$ (first canonical component) as the linear combination over the first dimension of matrices $\boldsymbol{A}$ and $\boldsymbol{B}$ that maximizes the Pearson correlation coefficient between the two vectors. Likewise, in PS analysis Korhonen and colleagues (Korhonen, Palva, and Palva, 2014) defined $g$ as the sparse weighted average over the first dimension that maximizes, in each area, the PS between the representative time series and all the sources in the area.

No matter how complex $g$ can be, all the approaches of the form share the fundamental property that function $h$, which estimates the FC, operates on the selected/constructed individual time series of each area (in the case of PS analysis, admittedly, including filtering, estimation of the HT and the phase of each time series). We, therefore, termed all these estimators of the form $f = h\big(g(\boldsymbol{A}, \boldsymbol{B})\big) = h(A, B)$ as *truly bivariate* or simply *bivariate*.

*Multivariate methods*

In the case of GPS/HPS, the function $f$ that estimates FC can again be decomposed, but differently:

$$\mathbb{R}^{N_A \times M} \times \mathbb{R}^{N_B \times M} \xrightarrow{g'} \mathbb{R}^{N_A \times N_B} \xrightarrow{h'} [0,1] \tag{6}$$

That is, here $f = h'\big(g'(\boldsymbol{A}, \boldsymbol{B})\big) = h'(\boldsymbol{PS})$, where $\boldsymbol{PS}$ is the pairwise inter-areal PS matrix whose elements are the pairwise values of the PS estimator between all the sources in areas $A$ and $B$. We termed this second approach as multivariate, although admittedly, the term mass-bivariate could also be a reasonable choice. Note that function $g'$ in (6) is practically identical to function $h$ in (5) (i.e., filtering, HT, phase extraction, and bivariate PS estimation). With the sole difference, of course, that the first three operations, which convert the raw time series into phase time series, are calculated $N_A + N_B$ times (one per each source in both areas), and the PS index is applied $N_A \times N_B$ times (all pair of sources in which each element belongs to a different area). Function $h'$ in (6) consists of simple algebraic operations such as averaging, as was the case of $g$ in (5).

In this case, by inverting the order of the operations, we eliminate the need to choose a proper way to reduce the dimensionality in the time domain, at the only (computational) cost of increasing the number of times PS is estimated (from 1 in (5) to $N_A \times N_B$ in (6)). More importantly, as we showed in the paper, we improved the consistency of the PS estimation.

It may be argued that a third family of methods that we may term *truly multivariate* will be that in which, after pre-processing (filtering, HT and phase estimation), PS is estimated among the $N_A$ and $N_B$ time series of phases in $A$ and $B$ without using any pairwise version or initial dimensionality reduction in the time domain. For instance, the distance covariance (Székely, Rizzo and Bakirov, 2007) is one of these genuinely multivariate approaches, which has been recently applied to fMRI analysis (Geerligs, Cam-CAN, and Henson, 2016), but its application in the context of PS synchronization has not been tested yet. The estimation of multivariate mutual information among the multivariate distribution of the two sets of phase series would fulfill such criteria. However, the estimation of multivariate marginal and joint entropies from this time series poses severe theoretical and practical difficulties (see, e.g. (Hlaváčková-Schindler *et al.*, 2007) for a review).

Therefore, in the present work, we compare the methods based on the approaches (5) and (6), using the same type of PS estimation but with an inversion in the order of application (on all sources or in the derived representative time series). Since the PS methods are the same, and the only difference is on which time series we apply them, it eases the interpretation of the results and the differences found between the two families of methods.

### Supplementary results

*Results for PLV-based metrics in all bands*

This work's main body shows the level of correlation between the FC estimated from EEG and MEG for the frequency bands theta, alpha, and low beta. We chose these frequency bands based on the rationale that both alpha and theta bands are expected to have a well-defined oscillator. In contrast, low beta is not (Bruña, Maestú, and Pereda, 2018), so that using these three bands would cover a broad range of possible cases. However, the correlation was calculated for all the classical bands: theta (4 to 8 Hz), alpha (8 to 12 Hz), low beta (12 to 20 Hz), high beta (20 to 30 Hz), and gamma (30 to 45 Hz). Delta band (2 to 4 Hz) was excluded from this study due to the high level of noise at these frequencies, usually rendering impossible the study of FC.

Supplementary Figure 3 shows the results for all five bands when using the classical (one-dimensional) procedure for PLV and ciPLV, and Supplementary Figure 4 shows the equivalent results obtained when using the three-dimensional procedure. As expected, the results for the high beta and gamma bands are better than for low beta and worse than for alpha and theta. This outcome is likely due to a poor fit of the real data in these bands to the PS model, even in the presence of oscillatory activity. Contrary to low beta, high beta and gamma are expected to manifest oscillators. However, both bands comprise a range of 10 Hz and are likely to contain more than one oscillator. Therefore, the phase extracted using Hilbert's transform will consist of a nonlinear mixture of several oscillators' phases. Also, EEG and MEG could present different sensitivities to different oscillators, so the corresponding weights in the mixture could differ, rising differences in the estimated phase and, with them, differences in the estimated FC.

Supplementary Table 1 shows the numerical results. As reported in the main text, the purely multivariate approaches are superior to that of the representative time series for all the metrics and bands, with the results for the approaches based on average PS and the RMS very similar and superior to those based on representative time series. We can also identify two groups of bands with different behaviors: in theta, alpha, high beta, and gamma bands, the results obtained using ciPLV are always superior to those obtained using PLV. However, in the low beta band, this effect is reversed, with the results obtained using PLV showing almost twice the number of significant correlations than those obtained using ciPLV. Still, even in the best case, the significant correlation ratio only reaches 18 % of the total number of comparisons, and none survives the FDR correction.

Supplementary Figure 3. Distribution of correlation coefficients between the multivariate PLV and ciPLV estimated using EEG and MEG for each approach and frequency band.

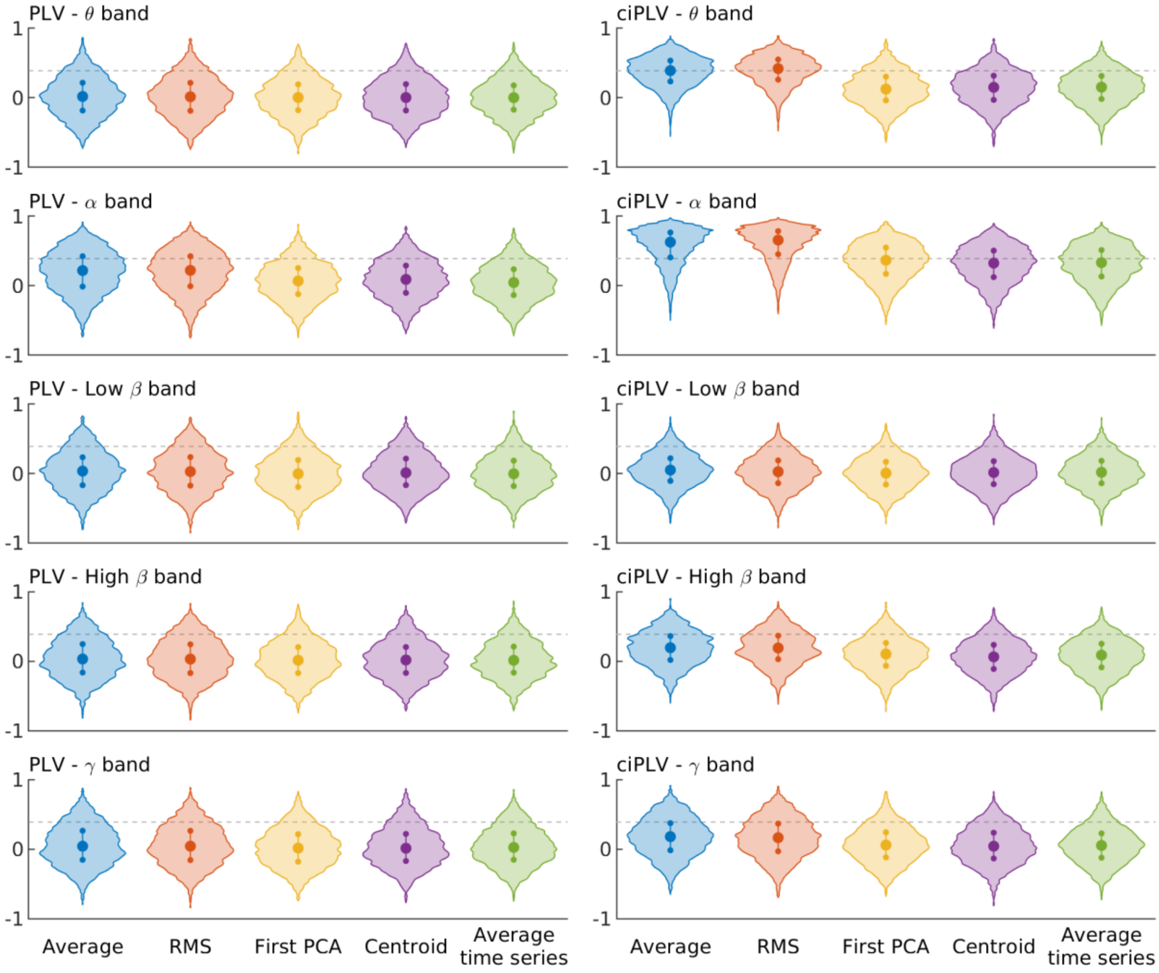


Violin plots showing the correlation between the inter-area PLV (left column) and ciPLV (right column) calculated with EEG and MEG using all the spatial dimensions and different approaches: Blue: Average of the pair-wise PS; Red: Root-mean-square (RMS) of the pair-wise PS; Yellow: PS between the PCA of each area; Purple: PS between the sources closest to the centroid of each area; Green: PS calculated the averaged time series of each area. Each row represents a classical frequency band. Dashed line marks the signification threshold with 5 % (one tailed) alpha level.

Supplementary Figure 4. Distribution of correlation coefficients between the multivariate PLV[3D] and ciPLV[3D] estimated using EEG and MEG for each approach and frequency band.

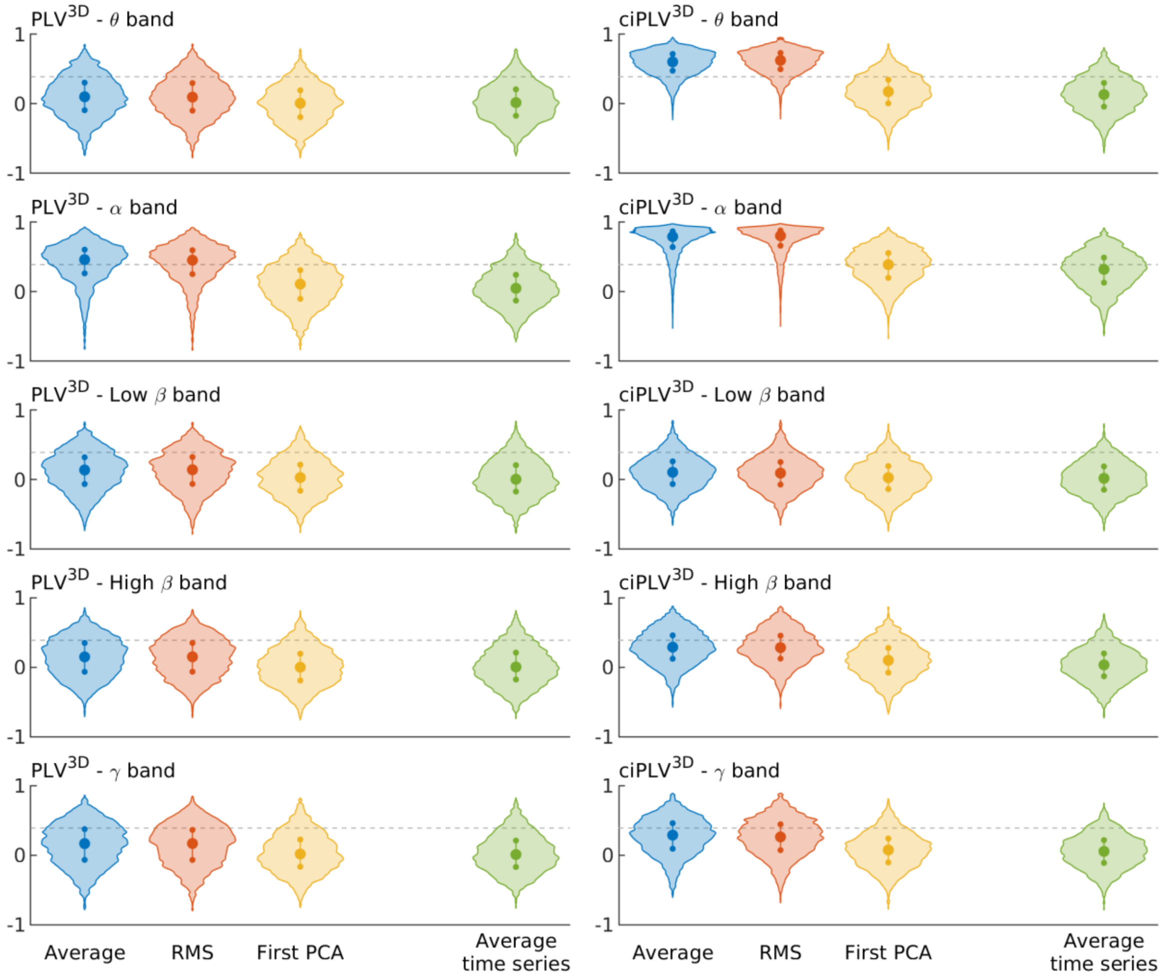


Violin plots showing the correlation between the inter-area PLV (left column) and ciPLV (right column) calculated with EEG and MEG using all the spatial dimensions and different approaches: Blue: Average of the pair-wise PS; Red: Root-mean-square (RMS) of the pair-wise PS; Yellow: PS between the PCA of each area; Green: PS calculated the averaged time series of each area. Each row represents a classical frequency band. Dashed line marks the signification threshold with 5 % (one tailed) alpha level.

Supplementary Table 1. Ratio of statistically significant correlation coefficients between the multivariate PS estimated using EEG and MEG for each approach and frequency band.

| | | Average PS | RMS | First PCA | Centroid | Average time series |
|---|---|---|---|---|---|---|
| PLV | Theta | 10 % (0 %) | 10 % (0 %) | 8 % (0 %) | 9 % (0 %) | 7 % (0 %) |
| | Alpha | 29 % (12 %) | 29 % (11 %) | 11 % (0 %) | 14 % (0 %) | 10 % (0 %) |
| | Low beta | 12 % (0 %) | 12 % (0 %) | 9 % (0 %) | 10 % (0 %) | 8 % (0 %) |
| | High beta | 12 % (0 %) | 12 % (0 %) | 8 % (0 %) | 9 % (0 %) | 9 % (0 %) |
| | Gamma | 14 % (1 %) | 13 % (0 %) | 10 % (0 %) | 11 % (1 %) | 11 % (0 %) |
| ciPLV | Theta | 50 % (32 %) | 56 % (38 %) | 15 % (0 %) | 15 % (0 %) | 16 % (0 %) |
| | Alpha | 76 % (74 %) | 80 % (78 %) | 47 % (32 %) | 40 % (23 %) | 42 % (25 %) |
| | Low beta | 8 % (0 %) | 6 % (0 %) | 5 % (0 %) | 6 % (0 %) | 6 % (0 %) |
| | High beta | 22 % (1 %) | 22 % (2 %) | 13 % (0 %) | 9 % (0 %) | 10 % (0 %) |
| | Gamma | 23 % (5 %) | 23 % (5 %) | 11 % (0 %) | 10 % (0 %) | 10 % (0 %) |
| $PLV^{3D}$ | Theta | 17 % (1 %) | 17 % (1 %) | 7 % (0 %) | - | 9 % (0 %) |
| | Alpha | 61 % (51 %) | 59 % (49 %) | 16 % (1 %) | - | 12 % (0 %) |
| | Low beta | 18 % (0 %) | 19 % (0 %) | 9 % (0 %) | - | 9 % (0 %) |
| | High beta | 20 % (1 %) | 20 % (0 %) | 9 % (0 %) | - | 9 % (0 %) |
| | Gamma | 23 % (4 %) | 22 % (1 %) | 11 % (0 %) | - | 10 % (0 %) |
| $ciPLV^{3D}$ | Theta | 87 % (85 %) | 89 % (88 %) | 20 % (1 %) | - | 15 % (0 %) |
| | Alpha | 91 % (90 %) | 91 % (91 %) | 50 % (35 %) | - | 40 % (21 %) |
| | Low beta | 11 % (0 %) | 10 % (0 %) | 7 % (0 %) | - | 6 % (0 %) |
| | High beta | 35 % (15 %) | 34 % (15 %) | 14 % (0 %) | - | 7 % (0 %) |
| | Gamma | 35 % (14 %) | 33 % (12 %) | 10 % (0 %) | - | 8 % (0 %) |

Ratio of statistically significant correlation coefficients (5 % alpha level, one tailed), and ratio of connections surviving false discovery rate (between brackets). PLV: Phase locking value. ciPLV: Corrected imaginary part of PLV. 3D superscript: PS calculated using all the spatial dimensions at each reconstructed source position. RMS: Root-mean-squared. PCA: Principal component analysis.

*Improvement when using all the meaningful orientations*

The direct observation of Supplementary Figure 3 and Supplementary Figure 4 suggests that the use of all the meaningful orientations at each source position improves the results for the purely multivariate approaches, but not necessarily for those based on representative time series. To get a more quantitative improvement indicator, we performed a paired t-test to compare the correlation coefficients' distribution using the one-dimensional (ci)PLV approach and the three-dimensional one. The results are depicted in Supplementary Table 2.

Supplementary Table 2. Comparison between the results attained using the one-dimensional source reconstruction and the three-dimensional source reconstruction for PLV and ciPLV for each approach and frequency band.

| | | **Average PS** | **RMS** | **First PCA** | **Average time series** |
|---|---|---|---|---|---|
| PLV | Theta | $p < 0.001$ | $p < 0.001$ | *n.s.* | *n.s.* |
| | Alpha | $p < 0.001$ | $p < 0.001$ | $p < 0.001$ | *n.s.* |
| | Low beta | $p < 0.001$ | $p < 0.001$ | $p = 0.002$ | *n.s.* |
| | High beta | $p < 0.001$ | $p < 0.001$ | *n.s.* | *n.s.* |
| | Gamma | $p < 0.001$ | $p < 0.001$ | *n.s.* | *n.s.* |
| ciPLV | Theta | $p < 0.001$ | $p < 0.001$ | $p < 0.001$ | *n.s.* |
| | Alpha | $p < 0.001$ | $p < 0.001$ | $p < 0.001$ | *n.s.* |
| | Low beta | $p < 0.001$ | $p < 0.001$ | $p = 0.001$ | *n.s.* |
| | High beta | $p < 0.001$ | $p < 0.001$ | *n.s.* | *n.s.* |
| | Gamma | $p < 0.001$ | $p < 0.001$ | *n.s.* | *n.s.* |

(Bonferroni-corrected) *p*-values for a (one tailed) t-test contrast of the distribution against the null hypothesis that the mean for the distribution using one- and three-dimensional procedures are the same. PLV: Phase locking value. ciPLV: Corrected imaginary part of PLV. 3D superscript: PS calculated using all the spatial dimensions at each reconstructed source position. RMS: Root-mean-squared. PCA: Principal component analysis. *n.s.*: Non-significant result ($p > 0.05$, one tailed).

The results show that the use of the three orientations improves those found for both multivariate approaches. They also improve for the representative time series based on the PCA, but only for some bands and metrics. Simultaneously, there is no significant improvement when using the average time series as a representative time series for the area. These results were expected since both purely multivariate approaches and (to a lesser extent) the PCA can successfully integrate large amounts of information. On the other hand, the increase of information could be detrimental to the average time series, as the likelihood of time series canceling out each other increases.

*Results for Hilbert coherence*

Throughout this work's main text, we have focused on estimating PS using PLV and ciPLV, as these metrics are extensively used in practice. However, we have also introduced a linear extension of PLV, namely Hilbert coherence (HCoh), which, although less extended in use, presents a series of advantages over PLV. To evaluate the ability of HCoh to estimate the degree of FC between cortical areas, we repeated the analysis using HCoh and its corrected imaginary counterpart (ciHCoh) instead of PLV and ciPLV. Supplementary Figure 5 and Supplementary Figure 6 depict the results for the one- and three-dimensional procedures, respectively, for each approach and frequency band.

Supplementary Figure 5. Distribution of correlation coefficients between the multivariate HCoh and ciHCoh estimated using EEG and MEG for each approach and frequency band.

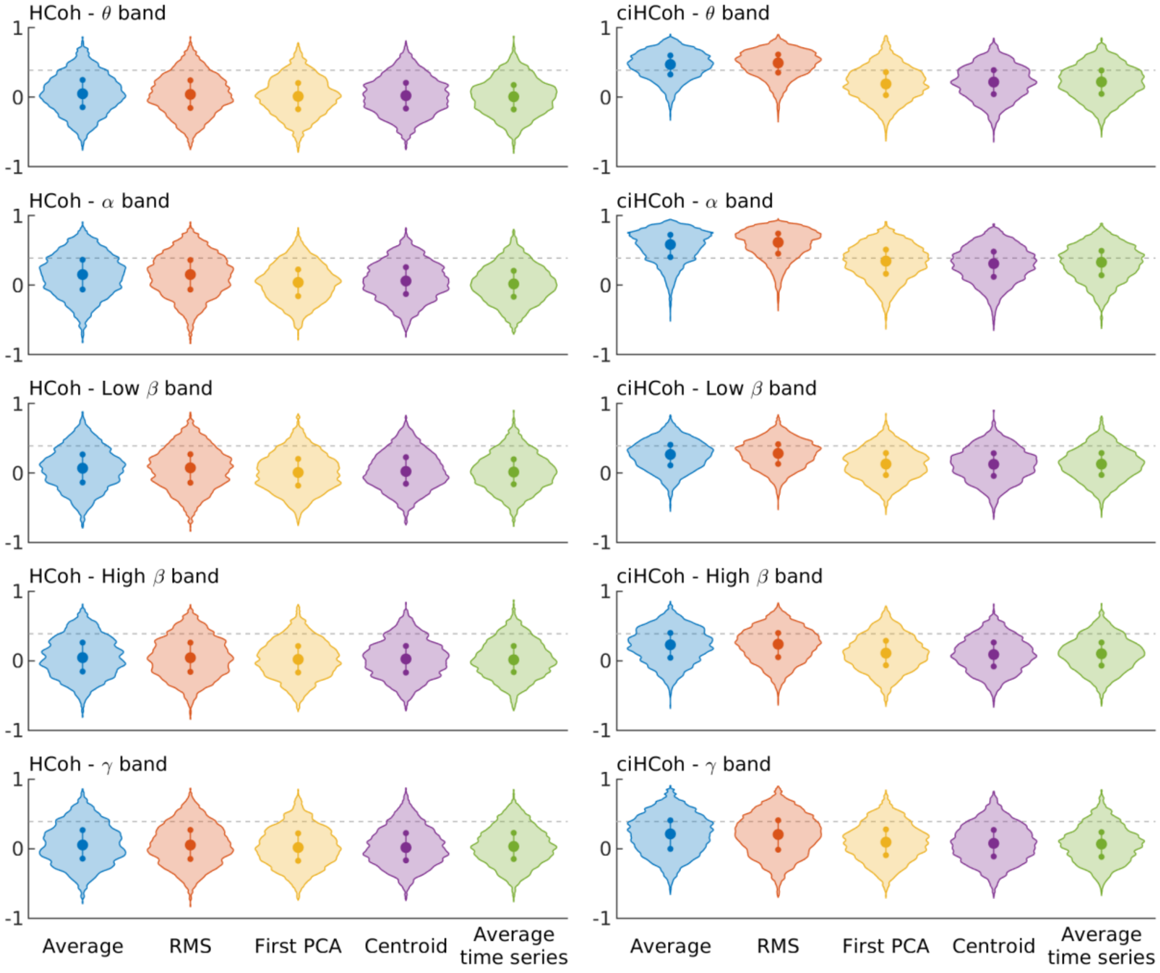


Violin plots showing the correlation between the inter-area HCoh (left column) and ciHCoh (right column) calculated with EEG and MEG using all the spatial dimensions and different approaches: Blue: Average of the pair-wise PS; Red: Root-mean-square (RMS) of the pair-wise PS; Yellow: PS between the PCA of each area; Purple: PS between the sources closest to the centroid of each area; Green: PS calculated the averaged time series of each area. Each row represents a classical frequency band. Dashed line marks the signification threshold with 5 % (one tailed) alpha level.

Supplementary Figure 6. Distribution of correlation coefficients between the multivariate HCoh$^{3D}$ and ciHCoh$^{3D}$ estimated using EEG and MEG for each approach and frequency band.

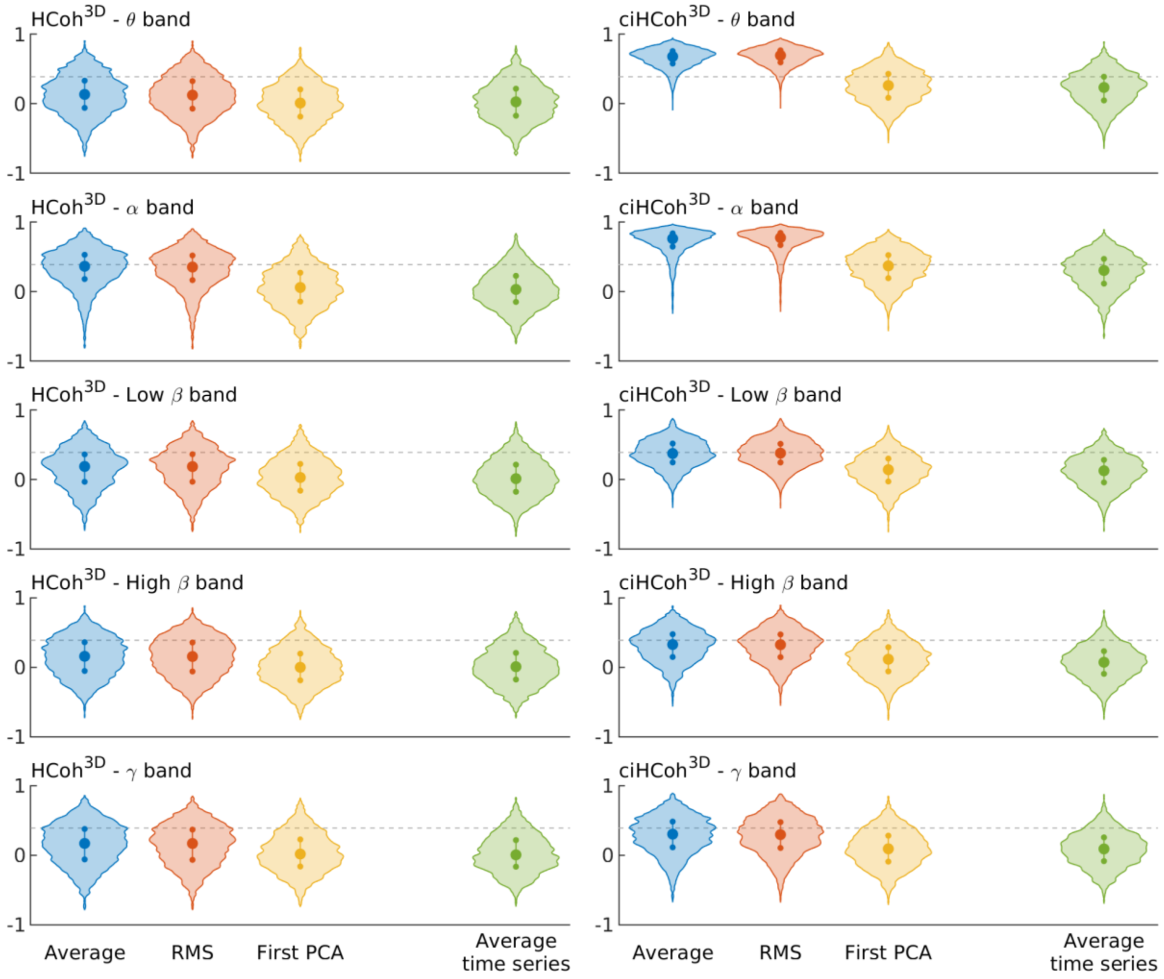


Violin plots showing the correlation between the inter-area HCoh (left column) and ciHCoh (right column) calculated with EEG and MEG using all the spatial dimensions and different approaches: Blue: Average of the pair-wise PS; Red: Root-mean-square (RMS) of the pair-wise PS; Yellow: PS between the PCA of each area; Green: PS calculated the averaged time series of each area. Each row represents a classical frequency band. Dashed line marks the signification threshold with 5 % (one tailed) alpha level.

Likewise, Supplementary Table 3 depicts the numerical results in terms of the ratio of significant correlation coefficients and the resulting distribution significance. The results are similar to those obtained with PLV and ciPLV, but with higher ratios of significant correlations. To test if this effect is significant, we calculated a (two-tailed) t-test comparing the results obtained with (ci)PLV and (ci)HCoh. The results are shown in Supplementary Table 4. For the purely multivariate approaches, there is a generalized improvement when using HCoh in theta, low beta, high beta, and gamma bands. However, this effect is reversed in the alpha band. The exception is ciPLV$^{3D}$ in the alpha band, where no differences between both metrics were found, likely because the results are packed in the higher extreme of the distribution.

Supplementary Table 3. Ratio of statistically significant correlation coefficients between the multivariate PS estimated using EEG and MEG for each approach and frequency band.

| | | Average PS | RMS | First PCA | Centroid | Average time series |
|---|---|---|---|---|---|---|
| HCoh | Theta | 12 % (0 %) | 11 % (0 %) | 8 % (0 %) | 9 % (0 %) | 7 % (0 %) |
| | Alpha | 22 % (5 %) | 22 % (5 %) | 10 % (0 %) | 12 % (0 %) | 9 % (0 %) |
| | Low beta | 14 % (1 %) | 14 % (1 %) | 9 % (0 %) | 11 % (0 %) | 8 % (0 %) |
| | High beta | 13 % (0 %) | 13 % (0 %) | 9 % (0 %) | 10 % (0 %) | 9 % (0 %) |
| | Gamma | 15 % (1 %) | 14 % (1 %) | 10 % (0 %) | 11 % (1 %) | 11 % (0 %) |
| ciHCoh | Theta | 66 % (55 %) | 70 % (61 %) | 21 % (4 %) | 25 % (2 %) | 25 % (2 %) |
| | Alpha | 76 % (73 %) | 82 % (80 %) | 43 % (25 %) | 38 % (19 %) | 41 % (21 %) |
| | Low beta | 28 % (1 %) | 29 % (1 %) | 13 % (0 %) | 13 % (0 %) | 13 % (0 %) |
| | High beta | 27 % (2 %) | 26 % (3 %) | 14 % (0 %) | 12 % (0 %) | 12 % (0 %) |
| | Gamma | 27 % (8 %) | 27 % (9 %) | 14 % (0 %) | 13 % (1 %) | 11 % (1 %) |
| $HCoh^{3D}$ | Theta | 20 % (3 %) | 19 % (2 %) | 9 % (0 %) | - | 10 % (0 %) |
| | Alpha | 46 % (30 %) | 45 % (27 %) | 13 % (1 %) | - | 11 % (0 %) |
| | Low beta | 22 % (3 %) | 22 % (3 %) | 10 % (0 %) | - | 9 % (0 %) |
| | High beta | 21 % (1 %) | 21 % (0 %) | 9 % (0 %) | - | 9 % (0 %) |
| | Gamma | 23 % (4 %) | 22 % (1 %) | 11 % (0 %) | - | 10 % (0 %) |
| $ciHCoh^{3D}$ | Theta | 95 % (95 %) | 97 % (97 %) | 31 % (7 %) | - | 25 % (2 %) |
| | Alpha | 93 % (93 %) | 94 % (94 %) | 47 % (29 %) | - | 38 % (16 %) |
| | Low beta | 47 % (26 %) | 48 % (25 %) | 14 % (0 %) | - | 12 % (0 %) |
| | High beta | 39 % (16 %) | 39 % (15 %) | 14 % (0 %) | - | 8 % (0 %) |
| | Gamma | 38 % (18 %) | 37 % (18 %) | 14 % (0 %) | - | 10 % (0 %) |

Ratio of statistically significant correlation coefficients (5 % alpha level, one tailed), and ratio of connections surviving false discovery rate (between brackets). HCoh: Hilbert coherence. ciHCoh: Corrected imaginary part of HCoh. 3D superscript: PS calculated using all the spatial dimensions at each reconstructed source position. RMS: Root-mean-squared. PCA: Principal component analysis.

The results for HCoh are very similar to those found with PLV, with the notable exception of ciHCoh in the low beta frequency band. While (ci)PLV was unable to detect any PS in this band, (ci)HCoh does, showing a level of agreement between EEG and MEG similar to that found in high beta or gamma bands. One possible reason for this result might be the nature of the low beta band's oscillatory activity. This band might be dominated by motor inhibitory activity, and this activity has been described as burst instead of a pure oscillatory rhythm (Jones, 2016). As PLV only considers the phase, the effect of the sparse bursts of beta activity would be negligible in the overall estimation of PS. However, as the bursts present a higher amplitude than the background beta activity, the amplitude weighting in HCoh enhances its influence in the estimated PS. This fact could be considered a weighting, depending on the signal to noise ratio of the signal, common to all coherence-based metrics (Bruña, Maestú, and Pereda, 2018).

Supplementary Table 4. Results for the comparison of performance of (ci)PLV and (ci)HCoh for each approach and frequency band.

| | | Average PS | RMS | First PCA | Centroid | Average time series |
|---|---|---|---|---|---|---|
| PLV vs. HCoh | Theta | $p < 0.001$ | $p < 0.001$ | $p < 0.001$ | $p < 0.001$ | *n.s.* |
| | Alpha | $p < 0.001$ | $p < 0.001$ | $p < 0.001$ | $p < 0.001$ | $p < 0.001$ |
| | Low beta | $p < 0.001$ | $p < 0.001$ | $p < 0.001$ | $p < 0.001$ | $p < 0.001$ |
| | High beta | $p < 0.001$ | $p < 0.001$ | $p < 0.001$ | $p = 0.001$ | *n.s.* |
| | Gamma | $p < 0.001$ | $p < 0.001$ | $p < 0.001$ | *n.s.* | $p < 0.001$ |
| ciPLV vs. ciHCoh | Theta | $p < 0.001$ | $p < 0.001$ | $p < 0.001$ | $p < 0.001$ | $p < 0.001$ |
| | Alpha | $p < 0.001$ | $p < 0.001$ | $p < 0.001$ | *n.s.* | *n.s.* |
| | Low beta | $p < 0.001$ | $p < 0.001$ | $p < 0.001$ | $p < 0.001$ | $p = 0.001$ |
| | High beta | $p < 0.001$ | $p < 0.001$ | *n.s.* | $p < 0.001$ | $p < 0.001$ |
| | Gamma | $p < 0.001$ | $p < 0.001$ | $p < 0.001$ | $p < 0.001$ | $p = 0.004$ |
| $PLV^{3D}$ vs. $HCoh^{3D}$ | Theta | $p < 0.001$ | $p < 0.001$ | $p < 0.001$ | - | $p < 0.001$ |
| | Alpha | $p < 0.001$ | $p < 0.001$ | $p < 0.001$ | - | $p < 0.001$ |
| | Low beta | $p < 0.001$ | $p < 0.001$ | $p < 0.001$ | - | $p = 0.017$ |
| | High beta | $p < 0.001$ | $p < 0.001$ | *n.s.* | - | *n.s.* |
| | Gamma | $p < 0.001$ | *n.s.* | *n.s.* | - | *n.s.* |
| $ciPLV^{3D}$ vs. $ciHCoh^{3D}$ | Theta | $p < 0.001$ | $p < 0.001$ | $p < 0.001$ | - | $p < 0.001$ |
| | Alpha | *n.s.* | *n.s.* | $p < 0.001$ | - | $p < 0.001$ |
| | Low beta | $p < 0.001$ | $p < 0.001$ | $p < 0.001$ | - | $p = 0.001$ |
| | High beta | $p < 0.001$ | $p < 0.001$ | $p = 0.004$ | - | $p < 0.001$ |
| | Gamma | $p < 0.001$ | $p < 0.001$ | $p = 0.001$ | - | $p < 0.001$ |

Bonferroni-corrected) *p*-values for a (two tailed) t-test contrast of the distribution against the null hypothesis that the mean of the distribution of (ci)PLV and (ci)HCoh is the same. PLV: Phase locking value. ciPLV: Corrected imaginary part of PLV. HCoh: Hilbert coherence. ciHCoh: Corrected imaginary part of HCoh. 3D superscript: PS calculated using all the spatial dimensions at each reconstructed source position. RMS: Root-mean-squared. PCA: Principal component analysis. *n.s.*: Non-significant result ($p > 0.05$, two tailed). The significant results show higher correlation coefficients for (ci)HCoh for all the frequency bands except for alpha, where the significant results show higher correlation coefficients for (ci)PLV.

When comparing the results obtained with PLV and HCoh (Supplementary Table 4), HCoh-based metrics are superior to PLV-based metrics in all the bands except alpha, where PLV is superior. This difference in the behavior of HCoh and PLV in the alpha band had already been observed (Bruña, Maestú, and Pereda, 2018). We hypothesize that it may be due to the different nature of the oscillations in this band. As the alpha oscillations are present almost always in resting-state data with eyes closed, weighting the signal by amplitude does not introduce an added benefit. This weighting could enhance the effect of high-amplitude artifacts, thus degrading the estimation's quality and explaining the superiority of PLV in this particular scenario.

*Results for identical relative bandwidth*

It can be argued that PS depends on the bandwidth, so a broader band entails an inherent penalty, as the phase variability is higher the broader the bandwidth. This fact could affect the results presented in the main text of this work, as the alpha band is narrower, in relative terms, than both theta and low beta. As defined here from 8 to 12 Hz, the relative bandwidth of alpha is 3:2, while for theta band (4 to 8 Hz), it is 2:1, and for low beta (12 to 20 Hz), it is 5:3. To explore this possibility, we repeated the analyses for the 3D versions of PLV and ciPLV using identical relative bandwidths, re-defining theta as 5 to 7.5 Hz and low beta as 13 to 19.5 Hz. These calculations are shown in Supplementary Figure 7 and Supplementary Table 5, where no clear improvement can be observed for the original data. In summary, while we can expect a penalty for broader bands when using phase synchronization, our results do not seem driven by it.

Supplementary Figure 7. Distribution of correlation coefficients between the multivariate $PLV^{3D}$ and $ciPLV^{3D}$ estimated using EEG and MEG for each approach and the re-defined frequency bands.

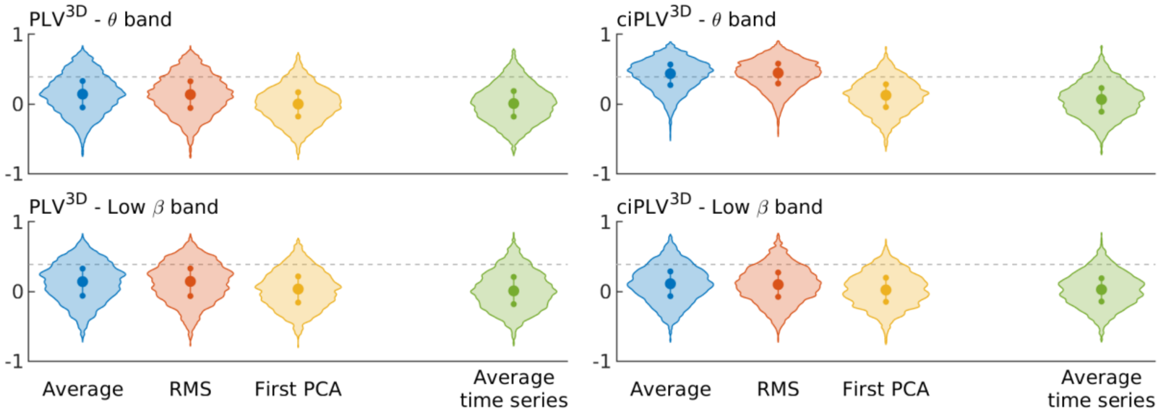


Violin plots showing the correlation between the inter-area $PLV^{3D}$ (left column) and $ciPLV^{3D}$ (right column) calculated with EEG and MEG using all the spatial dimensions and different approaches: Blue: Average of the pair-wise PS; Red: Root-mean-square (RMS) of the pair-wise PS; Yellow: PS between the PCA of each area; Green: PS calculated the averaged time series of each area. Each row represents a re-defined frequency band. Dashed line marks the signification threshold with 5 % (one tailed) alpha level.

Supplementary Table 5. Ratio of statistically significant correlation coefficients between the multivariate PS estimated using EEG and MEG for each approach and the re-defined frequency bands.

| | | **Average PS** | **RMS** | **First PCA** | **Centroid** | **Average time series** |
|---|---|---|---|---|---|---|
| $PLV^{3D}$ | Theta | 19 % (4 %) | 18 % (2 %) | 6 % (0 %) | - | 8 % (0 %) |
| | Low beta | 19 % (0 %) | 19 % (0 %) | 9 % (0 %) | - | 9 % (0 %) |
| $ciPLV^{3D}$ | Theta | 58 % (45 %) | 61 % (47 %) | 13 % (0 %) | - | 10 % (0 %) |
| | Low beta | 14 % (0 %) | 12 % (0 %) | 7 % (0 %) | - | 7 % (0 %) |

Ratio of statistically significant correlation coefficients (5 % alpha level, one tailed), and ratio of connections surviving false discovery rate (between brackets). PLV: Phase locking value. ciPLV: Corrected imaginary part of PLV. 3D superscript: PS calculated using all the spatial dimensions at each reconstructed source position. RMS: Root-mean-squared. PCA: Principal component analysis.

*Results using the average data for all the sessions*

All the results shown in the main text of this work and previous sections are based on using each session as an independent dataset. However, this might not be adequate, as 16 of the 19 sessions come from 8 participants (2 sessions each), and the remaining three sessions come from 3 participants. This fact could add some correlation between the 19 points used for the correlations and affect the results. To remove this possibility, we repeated the analyses using only 11 data points, using the average FC matrices for each participant. For those participants with two sessions, only one EEG and one MEG matrix were used, equal to the FC matrices' average from both sessions. For those participants with only one usable session, the FC matrices were unchanged. The results were similar to those found using the original 19 data points (this is, considering each session as an independent data). As an example of this, Supplementary Figure 8 shows the distribution of correlation coefficients for $PLV^{3D}$ and $ciPLV^{3D}$. The shape of the distribution shown here is similar to that in Supplementary Figure 4, but with a more strict threshold for significance (represented by the dashed line), due to the smaller sample size.

Supplementary Figure 8. Distribution of correlation coefficients between the multivariate $PLV^{3D}$ and $ciPLV^{3D}$ estimated using EEG and MEG for each approach and frequency band using only 11 data points (one per participant).

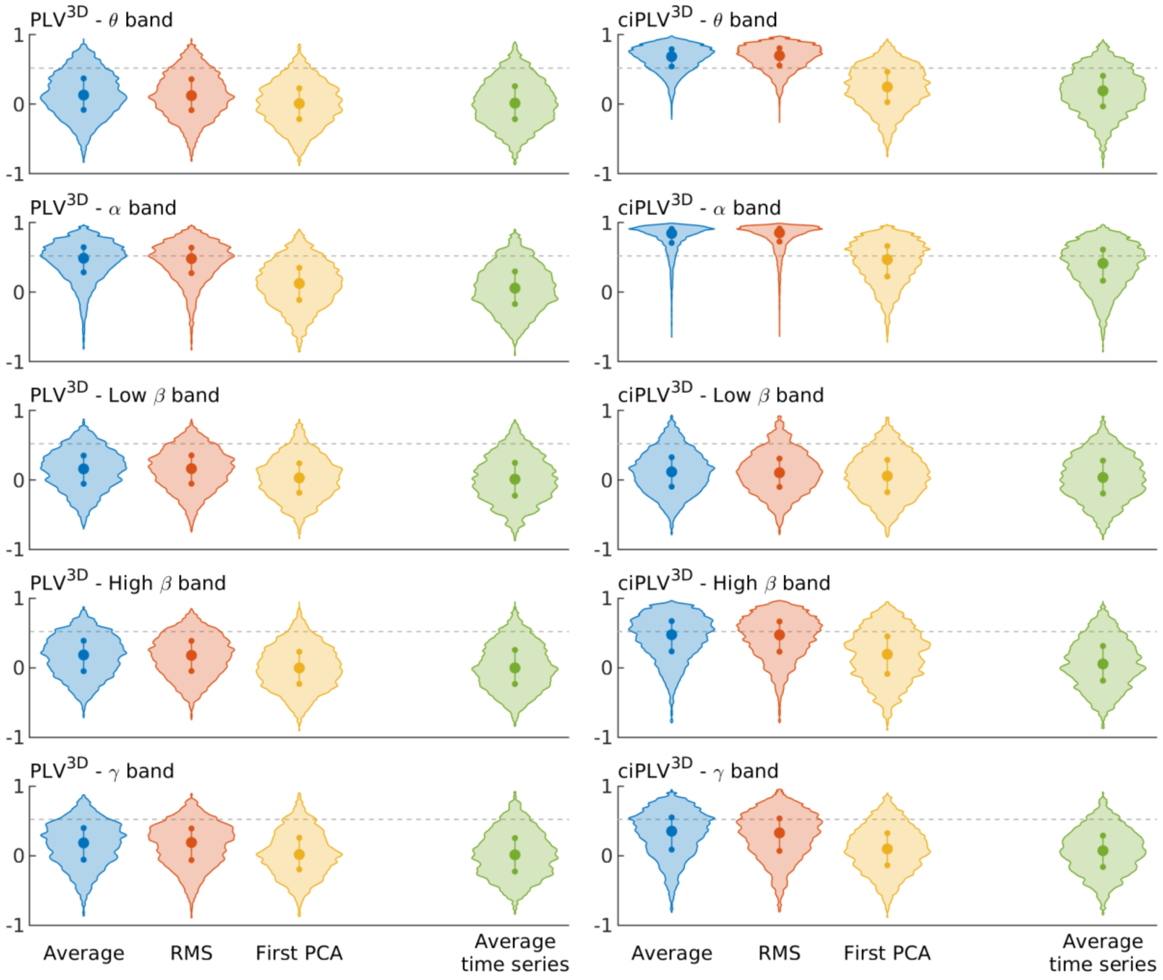


Violin plots showing the correlation between the inter-area PLV (left column) and ciPLV (right column) calculated with EEG and MEG using all the spatial dimensions and different approaches: Blue: Average of the pair-wise PS; Red: Root-mean-square (RMS) of the pair-wise PS; Yellow: PS between the PCA of each area; Green: PS calculated the averaged time series of each area. Each row represents a classical frequency band. Dashed line marks the signification threshold with 5 % (one tailed) alpha level.

Supplementary Table 6 and Supplementary Table 7 show the ratio of significant correlations for each metric and approach, both using an alpha level of 0.05 and using a false discovery rate. While the ratio of significant correlations is decreased for all cases, the relations between approaches are identical to the original scenario: the completely multivariate approaches outperform the representative time series.

Supplementary Table 6. Ratio of statistically significant correlation coefficients between the multivariate PS estimated using EEG and MEG for each approach and frequency band using only 11 data points (one per participant).

| | | **Average PS** | **RMS** | **First PCA** | **Centroid** | **Average time series** |
|---|---|---|---|---|---|---|
| PLV | Theta | 7 % (0 %) | 6 % (0 %) | 6 % (0 %) | 7 % (0 %) | 5 % (0 %) |
| | Alpha | 20 % (0 %) | 19 % (0 %) | 8 % (0 %) | 11 % (0 %) | 7 % (0 %) |
| | Low beta | 7 % (0 %) | 7 % (0 %) | 5 % (0 %) | 7 % (0 %) | 5 % (0 %) |
| | High beta | 7 % (0 %) | 7 % (0 %) | 5 % (0 %) | 7 % (0 %) | 7 % (0 %) |
| | Gamma | 8 % (0 %) | 8 % (0 %) | 6 % (0 %) | 8 % (1 %) | 8 % (0 %) |
| ciPLV | Theta | 42 % (10 %) | 45 % (15 %) | 15 % (0 %) | 14 % (0 %) | 15 % (0 %) |
| | Alpha | 71 % (67 %) | 74 % (70 %) | 43 % (23 %) | 37 % (15 %) | 38 % (17 %) |
| | Low beta | 8 % (0 %) | 6 % (0 %) | 6 % (0 %) | 6 % (0 %) | 7 % (0 %) |
| | High beta | 31 % (8 %) | 31 % (5 %) | 17 % (0 %) | 14 % (0 %) | 14 % (0 %) |
| | Gamma | 20 % (1 %) | 19 % (0 %) | 10 % (0 %) | 10 % (0 %) | 10 % (0 %) |
| $PLV^{3D}$ | Theta | 12 % (0 %) | 12 % (0 %) | 4 % (0 %) | - | 7 % (0 %) |
| | Alpha | 45 % (17 %) | 43 % (16 %) | 11 % (0 %) | - | 9 % (0 %) |
| | Low beta | 10 % (0 %) | 10 % (0 %) | 5 % (0 %) | - | 6 % (0 %) |
| | High beta | 12 % (0 %) | 12 % (0 %) | 5 % (0 %) | - | 7 % (0 %) |
| | Gamma | 12 % (0 %) | 11 % (0 %) | 7 % (0 %) | - | 7 % (0 %) |
| $ciPLV^{3D}$ | Theta | 78 % (72 %) | 81 % (75 %) | 19 % (0 %) | - | 15 % (0 %) |
| | Alpha | 89 % (88 %) | 90 % (90 %) | 44 % (15 %) | - | 36 % (13 %) |
| | Low beta | 9 % (0 %) | 8 % (0 %) | 8 % (0 %) | - | 7 % (0 %) |
| | High beta | 44 % (26 %) | 74 % (25 %) | 19 % (0 %) | - | 10 % (0 %) |
| | Gamma | 30 % (3 %) | 27 % (2 %) | 10 % (0 %) | - | 7 % (0 %) |

Ratio of statistically significant correlation coefficients (5 % alpha level, one tailed), and ratio of connections surviving false discovery rate (between brackets). PLV: Phase locking value. ciPLV: Corrected imaginary part of PLV. 3D superscript: PS calculated using all the spatial dimensions at each reconstructed source position. RMS: Root-mean-squared. PCA: Principal component analysis.

Supplementary Table 7. Ratio of statistically significant correlation coefficients between the multivariate PS estimated using EEG and MEG for each approach and frequency band using only 11 data points (one per participant).

| | | **Average PS** | **RMS** | **First PCA** | **Centroid** | **Average time series** |
|---|---|---|---|---|---|---|
| HCoh | Theta | 8 % (0 %) | 7 % (0 %) | 6 % (0 %) | 7 % (0 %) | 5 % (0 %) |
| | Alpha | 14 % (0 %) | 14 % (0 %) | 7 % (0 %) | 9 % (0 %) | 6 % (0 %) |
| | Low beta | 9 % (0 %) | 9 % (0 %) | 5 % (0 %) | 8 % (0 %) | 6 % (0 %) |
| | High beta | 7 % (0 %) | 7 % (0 %) | 5 % (0 %) | 7 % (0 %) | 7 % (0 %) |
| | Gamma | 8 % (0 %) | 8 % (0 %) | 6 % (0 %) | 8 % (1 %) | 8 % (0 %) |
| ciHCoh | Theta | 60 % (43 %) | 62 % (46 %) | 22 % (2 %) | 24 % (0 %) | 25 % (1 %) |
| | Alpha | 70 % (64 %) | 75 % (69 %) | 38 % (15 %) | 35 % (10 %) | 36 % (12 %) |
| | Low beta | 22 % (0 %) | 22 % (0 %) | 12 % (0 %) | 13 % (0 %) | 12 % (0 %) |
| | High beta | 32 % (6 %) | 32 % (7 %) | 19 % (0 %) | 16 % (0 %) | 16 % (0 %) |
| | Gamma | 24 % (1 %) | 23 % (1 %) | 12 % (0 %) | 12 % (0 %) | 11 % (0 %) |
| $HCoh^{3D}$ | Theta | 14 % (0 %) | 14 % (0 %) | 5 % (0 %) | - | 7 % (0 %) |
| | Alpha | 30 % (3 %) | 29 % (3 %) | 8 % (0 %) | - | 8 % (0 %) |
| | Low beta | 13 % (0 %) | 12 % (0 %) | 6 % (0 %) | - | 6 % (0 %) |
| | High beta | 11 % (0 %) | 11 % (0 %) | 5 % (0 %) | - | 6 % (0 %) |
| | Gamma | 11 % (0 %) | 11 % (0 %) | 7 % (0 %) | - | 7 % (0 %) |
| $ciHCoh^{3D}$ | Theta | 90 % (89 %) | 92 % (90 %) | 30 % (2 %) | - | 24 % (0 %) |
| | Alpha | 93 % (92 %) | 94 % (94 %) | 39 % (14 %) | - | 33 % (7 %) |
| | Low beta | 37 % (2 %) | 36 % (0 %) | 14 % (0 %) | - | 11 % (0 %) |
| | High beta | 42 % (21 %) | 42 % (20 %) | 20 % (1 %) | - | 11 % (0 %) |
| | Gamma | 31 % (4 %) | 29 % (4 %) | 12 % (0 %) | - | 9 % (0 %) |

Ratio of statistically significant correlation coefficients (5 % alpha level, one tailed), and ratio of connections surviving false discovery rate (between brackets). HCoh: Hilbert coherence. ciHCoh: Corrected imaginary part of HCoh. 3D superscript: PS calculated using all the spatial dimensions at each reconstructed source position. RMS: Root-mean-squared. PCA: Principal component analysis.

### References

Ahlfors, S. P. *et al.* (2010) 'Sensitivity of MEG and EEG to source orientation.', *Brain topography*, 23(3), pp. 227–32. doi: 10.1007/s10548-010-0154-x.

Basti, A. *et al.* (2018) 'Disclosing large-scale directed functional connections in MEG with the multivariate phase slope index', *NeuroImage*, 175, pp. 161–175. doi: 10.1016/j.neuroimage.2018.03.004.

Bruña, R., Maestú, F. and Pereda, E. (2018) 'Phase locking value revisited: Teaching new tricks to an old dog', *Journal of neural engineering*. Institute of Physics Publishing, 15(5), p. 056011. doi: 10.1088/1741-2552/aacfe4.

Dale, A. M. and Sereno, M. I. (1993) 'Improved localization of cortical activity by combining EEG and MEG with MRI cortical surface reconstruction: A linear approach', *Journal of Cognitive Neuroscience*, 5(2), pp. 162–176. doi: 10.1162/jocn.1993.5.2.162.

Ewald, A. *et al.* (2012) 'Estimating true brain connectivity from EEG/MEG data invariant to linear and static transformations in sensor space', *NeuroImage*, 60(1), pp. 476–488. doi: 10.1016/j.neuroimage.2011.11.084.

Fischl, B. R. *et al.* (2002) 'Whole brain segmentation: automated labeling of neuroanatomical structures in the human brain.', *Neuron*, 33(3), pp. 341–55.

Geerligs, L., Cam-CAN and Henson, R. N. (2016) 'Functional connectivity and structural covariance between regions of interest can be measured more accurately using multivariate distance correlation.', *NeuroImage*, 135, pp. 16–31. doi: 10.1016/j.neuroimage.2016.04.047.

Hlaváčková-Schindler, K. *et al.* (2007) 'Causality detection based on information-theoretic approaches in time series analysis', *Physics Reports*, 441(1), pp. 1–46. doi: 10.1016/j.physrep.2006.12.004.

Jones, S. R. (2016) 'When brain rhythms aren't "rhythmic": implication for their mechanisms and meaning', *Current Opinion in Neurobiology*, 40, pp. 72–80. doi: 10.1016/j.conb.2016.06.010.

Korhonen, O., Palva, S. and Palva, J. M. (2014) 'Sparse weightings for collapsing inverse solutions to cortical parcellations optimize M/EEG source reconstruction accuracy', *Journal of Neuroscience Methods*, 226, pp. 147–160. doi: 10.1016/j.jneumeth.2014.01.031.

Murakami, S. and Okada, Y. (2006)' Contributions of principal neocortical neurons to magnetoencephalography and electroencephalography signals', *The Journal of physiology*, 575(3), pp. 925–936. doi: 10.1113/jphysiol.2006.105379.

Sarvas, J. (1987) 'Basic mathematical and electromagnetic concepts of the biomagnetic inverse problem.', *Physics in Medicine and Biology*. IOP Publishing, 32(1), pp. 11–22. doi: 10.1088/0031-9155/32/1/004.

Székely, G. J., Rizzo, M. L. and Bakirov, N. K. (2007) 'Measuring and testing dependence by correlation of distances', *The Annals of Statistics*, 35(6), pp. 2769–2794. doi: 10.1214/009053607000000505.

Vrba, J. and Robinson, S. E. (2001) 'Signal processing in magnetoencephalography.', *Methods (San Diego, Calif.)*, 25, pp. 249–271. doi: 10.1006/meth.2001.1238.
